# Neuronal glutamate transporters control dopaminergic signaling and compulsive behaviors

**DOI:** 10.1101/224477

**Authors:** Stefania Bellini, Kelsey E. Fleming, Modhurika De, John P. McCauley, Maurice A. Petroccione, Lianna Y. D’Brant, Artem kachenko, SoYoung Kwon, Lindsey A. Jones, Annalisa Scimemi

**Affiliations:** SUNY Albany, Department of Biology, 1400 Washington Avenue, Albany (NY) 12222, USA

**Author notes:** These authors contributed equally to this work (Stefania Bellini, Kelsey E. Fleming, Modhurika De). **CORRESPONDING AUTHOR** Dr. Annalisa Scimemi, PhD SUNY Albany, Department of Biology, 1400 Washington Avenue, Albany (NY) 12222, USA.

## Abstract

There is an ongoing debate on the contribution of the neuronal glutamate transporter EAAC1 to the onset of compulsive behaviors. Here we use behavioral, electrophysiological, molecular and viral approaches in male and female mice to identify the molecular and cellular mechanisms by which EAAC1 controls the execution of repeated motor behaviors. Our findings show that in the striatum, a brain region implicated with movement execution, EAAC1 limits group I metabotropic glutamate receptor (mGluRI) activation, facilitates D1 dopamine receptor (D1R) expression and ensures long-term synaptic plasticity. Blocking mGluRI in slices from mice lacking EAAC1 restores D1R expression and synaptic plasticity. Conversely, activation of intracellular signaling pathways coupled to mGluRI in D1R-expressing striatal neurons of mice expressing EAAC1 leads to reduced D1R expression and increased stereotyped movement execution. These findings identify new molecular mechanisms by which EAAC1 can shape glutamatergic and dopaminergic signals and control repeated movement execution.

**SIGNIFICANCE STATEMENT:** Genetic studies implicate *Slc1a1*, a gene encoding the neuronal glutamate transporter EAAC1, with obsessive-compulsive disorder (OCD). EAAC1 is abundantly expressed in the striatum, a brain region that is hyperactive in OCD. What remains unknown is how EAAC1 shapes synaptic function in the striatum. Our findings show that EAAC1 limits activation of metabotropic glutamate receptors (mGluRI) in the striatum and, by doing so, it promotes D1R expression. Targeted activation of signaling cascades coupled to mGluRI in mice expressing EAAC1 reduces D1R expression and triggers repeated motor behaviors in mice. These findings provide new information on the molecular basis of OCD and suggest new avenues for its treatment.

## INTRODUCTION

Persistent thoughts, anxiety and repeated execution of stereotyped movements are hallmark features of neuropsychiatric disorders like OCD (Kessler et al., 2005). Family-based linkage analysis identify the gene *Slc1a1*, which encodes the neuronal glutamate transporter EAAC1, as one of the strongest candidate genes for OCD (Hanna et al., 2002; Arnold et al., 2006; Dickel et al., 2006; Stewart et al., 2007; Shugart et al., 2009; Wendland et al., 2009; Samuels et al., 2011). Alternative isoforms of *Slc1a1* that are differentially regulated in OCD patients have been shown to impair glutamate uptake via EAAC1 (Porton et al., 2013). One of the hypotheses that has been put forward is that loss of function of EAAC1 leads to increased extracellular glutamate concentration and hyperactivity in the brain (Porton et al., 2013). This hypothesis is not consistent with functional studies *in vitro* indicating that the regulation of the ambient glutamate concentration in the brain does not rely on *neuronal* but on *glial* glutamate transporters (Jabaudon et al., 1999; Cavelier and Attwell, 2005; Le Meur et al., 2007), which are hundred times more abundantly expressed than neuronal transporters (Holmseth et al., 2012). Our own previous work shows that EAAC1 exerts a powerful control of phasic glutamatergic synaptic transmission but does not alter the ambient glutamate concentration in the hippocampus (Scimemi et al., 2009). Despite these findings, we still have limited knowledge on the function of EAAC1 in other regions of the brain that show structural and functional abnormalities in patients with OCD, like the striatum.

The striatum is the main entry point of excitatory inputs to the basal ganglia and exerts a fundamental role in the control of anxiety and movement execution (Piras et al., 2015). In the striatum, glial glutamate transporters clear synaptically-released glutamate from the extracellular space and regulate GluA/N activation, similar to what they do in the hippocampus (Goubard et al., 2011). EAAC1 is abundantly expressed in the striatum, but its role in regulating synaptic function has so far remained elusive (Danbolt, 2001; Holmseth et al., 2012). Here we show that loss of EAAC1 is associated with increased stereotyped movement execution and anxiety-like behaviors in mice. In the striatum, EAAC1 limits mGluRI activation and, by doing so, it promotes D1R expression. Blocking mGluRI in mice lacking EAAC1 and cell-specific activation of signaling cascades coupled to mGluRI in mice expressing EAAC1 allow for bi-directional control of D1R expression, synaptic plasticity and repeated movement execution. These results identify new molecular mechanisms by which EAAC1 controls the function of the striatum and point to its pivotal role as a molecular switch to control mGluRI activation, glutamatergic and dopaminergic transmission, and ultimately the execution of persistent motor behaviors.

## MATERIALS AND METHODS

### Ethics statement

All experimental procedures were performed in accordance with protocols approved by the Institutional Animal Care and Use Committee at SUNY Albany and guidelines described in the US National Institutes of Health *Guide for the Care and Use of Laboratory Animals*.

### Mice

All mice were group-housed and kept under a 12 hour light cycle (6:00 AM ON, 6:00 PM OFF) with food and water available *ad libitum*. Constitutive EAAC1 knock-out mice (EAAC1^−/−^) were obtained by targeted disruption of the *Slc1a1* gene via insertion of a pgk neomycin resistance cassette (Neo) in exon 1 of the *Slc1a1* gene, as originally described by (Peghini et al., 1997). EAAC1^−/−^ breeders were generated after back-crossing EAAC1^+/−^ mice with C57BL/6 mice for more than 10 generations, as described by (Scimemi et al., 2009). C57BL/6 wild-type (WT) and EAAC1^−/−^ mice (P0-35) were identified by PCR analysis of genomic DNA. EAAC1^−/−^ mice develop normally during the first five weeks of post-natal life. They are fertile and although they give birth to smaller litters (number of pups in each litter: WT 8.2±0.3 (n=43), EAAC1^−/−^ 6.5±0.4 (n=42), ***p=8.0e-4), the litters are as viable as those of WT mice (peri-natal mortality rate: WT 0.23±0.04 (n=40), EAAC1^−/−^ 0.30±0.06 (n=41), p=0.30) and have a similar sex distribution (proportion of females in each litter: WT 0.47±0.04 (n=32), EAAC1^−/−^ 0.51±0.03 (n=26), p=0.52). These data are consistent with previous phenotypic characterization of EAAC1^−/−^ mice (Peghini et al., 1997).

D1^Cre/+^ mice (MMRRC Cat# 030778-UCD; STOCK Tg(Drd1-cre)EY217Gsat/Mmucd) and A2A^Cre/+^ mice (MMRRC Cat# 036158 B6.FVB(Cg)-Tg(Adora2a-cre)KG139Gsat/Mmucd) (Gong et al., 2003; Gong et al., 2007) were kindly provided by Drs. A.V. Kravitz and C.F. Gerfen (NIH/NIDDK). In these mice, the protein Cre-recombinase is expressed under the control of the promoter for D1Rs and the adenosine receptor 2 (which co-localizes with D2 dopamine receptors (D2Rs)) respectively. Ai9^Tg/Tg^ conditional reporter mice (The Jackson Laboratory Cat# 007909; B6.Cg-Gt(ROSA)26Sor^tm9(CAG-tdTomato)Hze^) (Madisen et al., 2010) were kindly provided by Dr. P.E. Forni (SUNY Albany). D1^tdTomato/+^ mice were purchased from The Jackson Laboratory (Cat# 016204; Tg(Drd1a-tdTomato)6Calak).

Genotyping was performed on toe tissue samples at P7-10. Briefly, tissue samples were digested at 55°C overnight in a lysis buffer containing (mM): Tris base pH 8 (100), EDTA (5), NaCl (200), 0.2% SDS, and 100 μg/ml Proteinase K. DNA samples were diluted in nuclease-free water (500 ng/μl) and processed for PCR analysis. The PCR primers used for EAAC1, D1R^Cre/+^, A2A^Cre/+^, D1td^Tomato/+^ and Ai9 were purchased from Fisher Scientific (Hampton, NH) and their nucleotide sequence is listed in **Table 1.** The PCR protocol for EAAC1, D1R^Cre/+^, A2A^Cre/+^, D1^tdTomato/+^ and Ai9 are described in **Table 2-5.** PCR reactions for D1^tdTomato/+^ were performed using a Hotstart Taq polymerase (Cat# KK5621; KAPA Biosystems, Wilmington, MA). For all other reactions we used standard Taq DNA Polymerase (Cat# R2523, Millipore Sigma, St.Louis, MO).

**Table 1.**
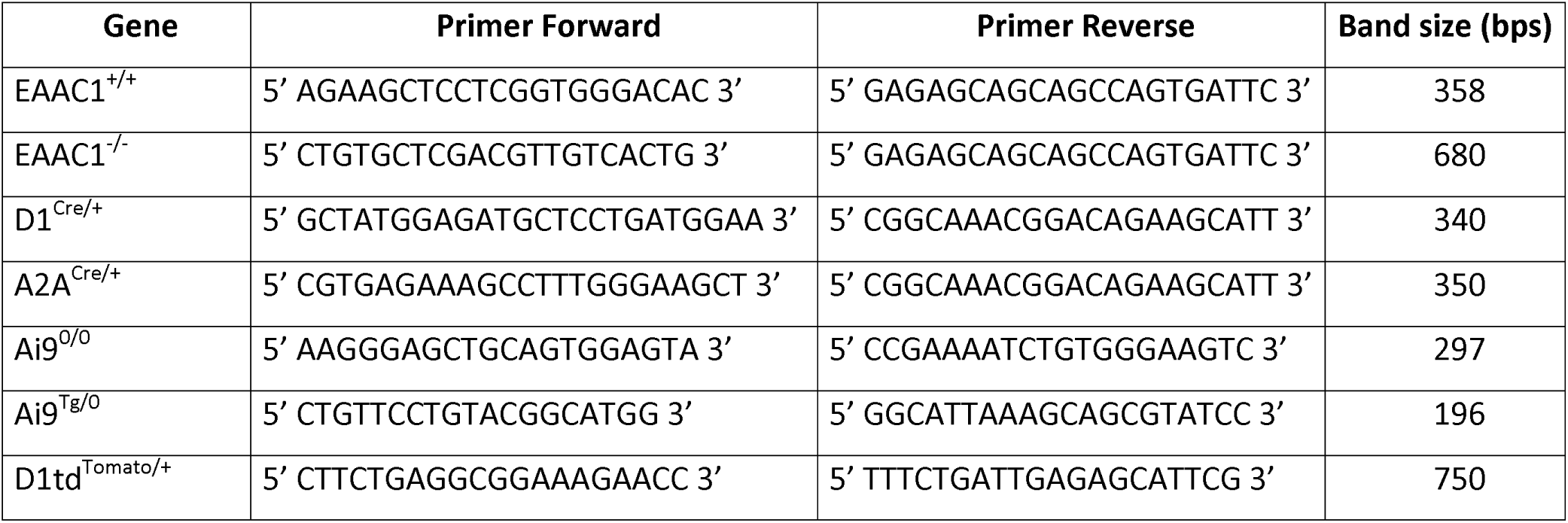
Sequence of primers used for PCR analysis.

**Table 2.**
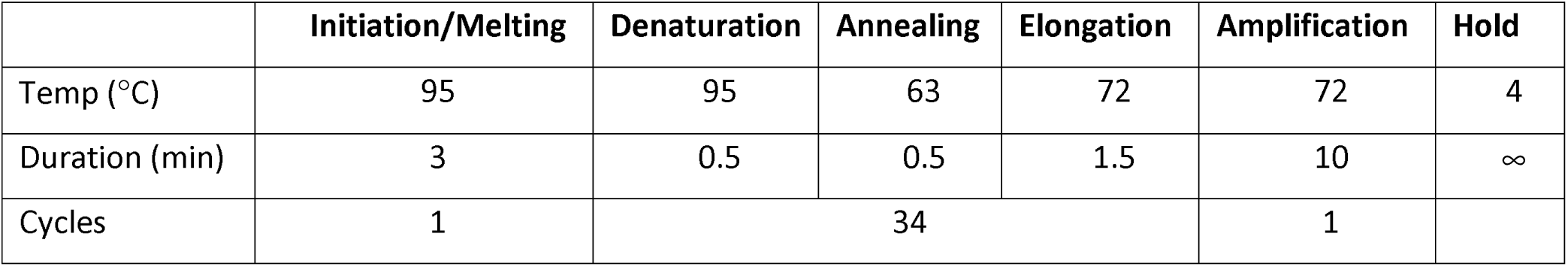
PCR protocol for EAAC1.

**Table 3.**
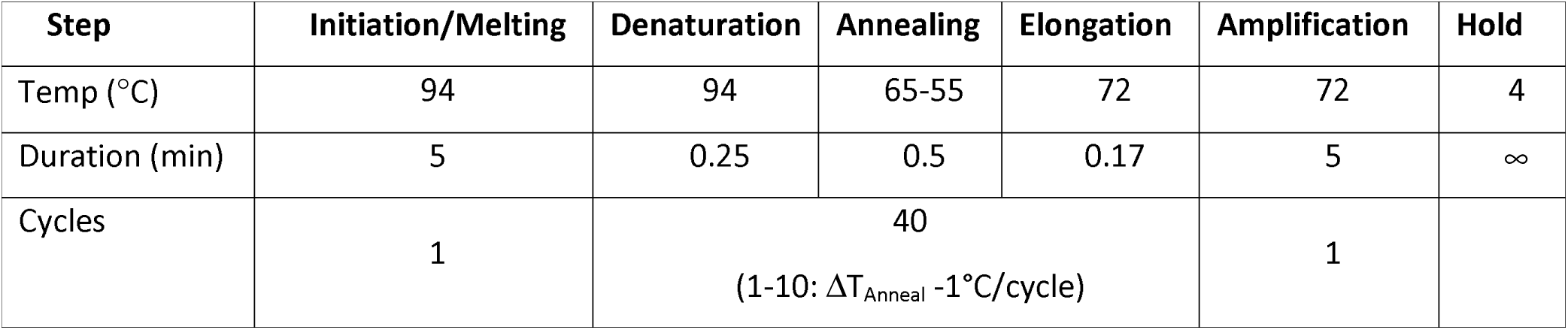
PCR protocol for D1^Cre/+^and A2A^Cre/+^.

**Table 4.** PCR protocol for D1'^dTomato/+^.

| Step | Initiation/Melting | Denaturation | Annealing | Elongation | Amplification | Hold |
| --- | --- | --- | --- | --- | --- | --- |
| Temp (°C) | 95 | 95 | 60 | 72 | 72 | 4 |
| Duration (min) | 3 | 0.25 | 0.25 | 0.25 | 1 | ∞ |
| Cycles | 1 | 35 |  |  | 1 |  |

**Table 5.** PCR protocol for Ai9.

| Step | Initiation/Melting | Denaturation | Annealing | Elongation | Amplification | Hold |
| --- | --- | --- | --- | --- | --- | --- |
| Temp (°C) | 94 | 94 | 65-60 | 72 | 72 | 10 |
| Duration (min) | 2 | 0.33 | 0.25 | 0.17 | 2 | ∞ |
| Cycles | 1 | 38<br>(1-10: $\Delta T_{\text{Anneal}} -0.5^{\circ}\text{C}/\text{cycle}$) | | | 1 | |

### Behavior

Before performing any behavioral test, mice were acclimated to a new behavioral suite for at least 30 min. All mice were tested between 9:00 AM and 1:00 PM. A battery of behavioral tests was performed on naïve WT and EAAC1^−/−^ mice (P14-35) and on WT and D1^Cre/+^ and A2A^Cre/+^ mice subjected to AAV-hSyn-DIO-hM3D(Gq)-mCherry stereotaxic and CNO I/P injections, according to the experiments included in the Results section of the paper. The battery of behavioral tests included grooming, SmithKline-Beecham Harwell Imperial-College and Royal-London-Hospital Phenotype Assessment (SHIRPA), flying saucer, open field, elevated plus maze and marble burying tests, performed as described below. All behavioral apparatus were cleaned with 70% ethanol between each test.

### Grooming test

The behavioral arena used to acquire videos for the grooming analysis consisted of four chambers with clear bottom and white side walls (L 15 cm x W 25 cm x H 15 cm). A digital SLR camera (Canon EOS Rebel T3i with EF-S 18-55 mm f/3.5-5.6 IS lens, 60 fps) was positioned 6-12” below the grooming chamber to acquire 10 min long videos in which we monitored the grooming behavior of each mouse. The distance travelled by the forepaws during grooming was analyzed using a newly-developed tracking algorithm (Reeves et al., 2016). The identification of the phase composition of each grooming episode was performed according to the scaling system described by (Kalueff et al., 2007). Briefly, we identified no grooming (Phase 0), paw licking (Phase 1), face wash (Phase 2), body grooming (Phase 3), hind leg licking (Phase 4) and tail/genitals grooming (Phase 5). Correct transitions between grooming phases include the progressive transitions through all the steps of a grooming syntactic chain (e.g. 0-1, 1-2, 2-3, 3-4, 4-5 and 5-0). Any other transition that did not follow this order was classified as incorrect (Kalueff et al., 2007).

### SHIRPA test

The SHIRPA protocol is a collection of simple tests that we used to provide a standardized, high-throughput screen for assessing the phenotype of WT and EAAC1^−/−^ mice (P14-35) (Rogers et al., 1997; Rogers et al., 1999). The SHIRPA test is effective in distinguishing qualitative differences between different strains of mice. The following tests and scores were included in the SHIRPA protocol.

1. Condition score: (1-5). Emaciated (1), thin (2), normal (3), over conditioned (4), obese (5).
2. Gait: (0-1). Monitors exaggerated limb movements, dragging and uneven cadence. Abnormal (0), normal (1).
3. Posture: (0-1). Monitors the presence of rounded or hunched body, head tilt and tail dragging. Abnormal (0), normal (1).
4. Body tone: (0-3). Hold the mouse by the tail base on a hard surface and gently press with two fingers over the mid dorsum. Flaccid (0), allows depression to the floor (1), allows some flattening (2), hunches back to completely resist compression (3).
5. Petting escape: (0-3). Hold the mouse by the tail base on a hard surface and stroke down its flanks from front to back. No reaction (0), difficult to elicit escape response (1), easy to elicit escape response (2), difficult to test because of spontaneous escape attempts (3).
6. Passivity: (0-3). Hold the mouse by the tail and place the front paws on the edge of the cage top. Normal mice promptly climb up to the top of the cage. Falling off or hanging without climbing is abnormal. Falls (0), delayed or unsuccessful attempt to climb up (1), normal (2), hyperactive (3).
7. Trunk curl: (0-3). Suspend the mouse from the tail for 15 s and monitor for curling of trunk. Zero or abnormal response (e.g. hindlimb clasping) (0), curls <90° (1), curls to 90° or more (2), climbs up the tail (3).
8. Righting: (0-3). Use one hand to hold the mouse by the tail base and the other hand to provide a vertical surface. Normal mice feel the surface of the hand and quickly flip over. Mouse does not right itself (0), struggles to right itself (1), rights itself (2), hyperactive (3).
9. Visual placing/Reach touch: (0-1). Hold the mouse by the tail and lower it slowly toward the cage lid. Blind mice do not reach out until forelimbs or whiskers touch (0). Normal mice start to reach towards the surface well before touching it (1).
10. Whisker response: (0-3). Stimulate the vibrissae using a cotton tipped applicator. Touching vibrissae should elicit a response, including cessation of “whisking” or a subtle responsive nose quiver. No response (0), response difficult to elicit (1), normal response (2), hyperactive response (3).
11. Ear twitch: (0-3). Use a cotton tipped applicator to gently touch the ear pinna. A normal response is a rapid ear twitch. No response (0), difficult to elicit response (1), obvious response (2), hyper repetitive response (3).
12. 1Palpebral reflex: (0-3). Use a cotton tipped applicator to gently touch the cornea. No reaction (0), slow blink (1), quick blink (2), hyper repetitive blinking (3).
13. Forelimb place: (0-3). Hold the mouse by the tail on a hard surface and, using a cotton tipped applicator, we gently move a forelimb out to the side. A normal response is to immediately return the limb under the body. Leg stays where placed (0), slow or incomplete return (1), promptly returns the leg to normal position (2), hyperactive response (3).
14. 14. Withdrawal: (0-3). Hold the mouse by the tail on a hard surface, pick up the hind-limb and pull the limb out at a 45° angle until it is stretched and then let go. A normal mouse rapidly returns the hind-limb to normal position. Leg drops to ground and doesn’t return to normal position (0), leg slow to return (1), rapid return (2), hyperactive response (3).
15. Biting: (0-1). Place a wooden stick in front of the mouse’s mouth. The most common reaction is to ignore or turn away from the stick. No biting (0), biting (1).
16. Clicker (hearing test): (0-3). Hold the mouse by the tail base on a hard surface and use a clicker once after a moment of silence, to monitor for an ear flick or stop response. No response (0), difficulty in eliciting response (1), immediate response (2), abnormal response (3).
17. Grip: (s). Place a mouse on a wire metal grid 1-2' above ground, start a timer, shake the grid gently, rapidly flip it over and measure the time until the mouse falls off the grid.

### Flying saucer (running wheel) test

A plastic flying saucer disk (∅=5.25”) connected to an odometer (Model# SD-548B, Shenzhen Sunding Electron Co., Shenzhen, China) was positioned in a standard rat cage. The distance and time spent on the flying saucer were monitored over a period of 30 min.

### Open field test

In the open field test, we monitored the position of a mouse freely moving in a white Plexiglas box (L 46 cm × W 46 cm × H 38 cm). Each mouse was video monitored for 15 min using a Live! Cam Sync HD webcam (Model# VF0770; Creative Labs, Milpitas, CA). Videos were analyzed using AnyMaze (Stoelting, Wood Dale, IL).

### Elevated plus maze test

The elevated plus maze consisted of two open and two closed arms (L 35.6 cm x W 5 cm) that extended from a center platform (L 5 cm × W 5 cm) elevated 52 cm from the floor. Each mouse was placed in the center area of the elevated plus maze facing an open arm and allowed to move freely between the arms for 15 min. Each mouse was video monitored for 15 min using a Live! Cam Sync HD webcam (Model# VF0770; Creative Labs, Milpitas, CA). Videos were analyzed using AnyMaze (Stoelting, Wood Dale, IL). The number of entries and the amount of time spent in the open and closed arms were assessed as indices of anxiety-like behaviors.

### Marble-burying test

We filled a mouse cage with 5 cm bedding material and on top of it we arranged 24 glass marbles (∅=0.6 cm) in a 4 × 6 grid (distance from cage walls=1.3 cm; distance between marbles=3.8 cm). Each mouse spent 30 min in this cage. At the end of this time, we counted the number of marbles that had ≥50% of their top surface covered with bedding material.

## Real-time quantitative PCR

We prepared acute brain slices from WT and EAAC1^−/−^ mice of either sex (P16-24), deeply anesthetized with isoflurane and decapitated in accordance with SUNY Albany Animal Care and Use Committee guidelines. The brain was rapidly removed and placed in ice-cold slicing solution bubbled with 95% 0_2^-^_5% CO_2_, containing (in mM): 119 NaCl, 2.5 KCl, 0.5 CaCl_2_, 1.3 MgSO_4_·H_2_O, 4 MgCl_2_, 26.2 NaHCO_3_, 1 NaH_2_PO_4_, and 22 glucose; 320 mOsm; pH 7.4. Coronal brain slices (250 μm thick) were prepared using a vibrating blade microtome (VT1200S, Leica Microsystems, Wetzlar, Germany). The slices were transferred to an ice-cold RNA stabilizing solution (Cat# 76106, Qiagen, Valencia, CA) in which we separated the dorsolateral (DLS) from the ventromedial striatum (VMS). The total RNA was purified and transcribed into cDNA using Taq DNA polymerase (Cat# 201205, Qiagen, Valencia, CA) according to the manufacturer’s instructions. We measured transcriptional levels of *Drd1a* and *Drd2* in triplicate samples using the TaqMan gene expression assays (Thermo Fisher Scientific, Waltham, MA). Each PCR reaction was run as a duplex reaction using the housekeeping gene *Hprt1* as an internal control. We used the following primers to measure the amount of cDNA for *Drd1a, Drd2* and *Hprt: Drd1a* (Mm02620146_s1, FAM-MGB), *Drd2* (Mm00438545_m1, FAM-MGB), *Slc1a1* (Mm00436587_m1, FAM-MGB), *Hprt1* (Mm00446968_m1, VIC-MGB) (Thermo Fisher Scientific, Waltham, MA). All reactions were performed using either the TaqMan Universal Master Mix II (Thermo Fisher Scientific, Waltham, MA) or nuclease-free water (Millipore Sigma, St. Louis, MO). Initial validation experiments were performed using a range of cDNA dilutions, to ensure similar amplification efficiency of all target cDNAs. A no template control (NTC), in which cDNA was omitted from the reaction mix, was used to monitor contamination and primer-dimer formation that could produce false positive results. In each experiment, the threshold cycle (C_T_) at which fluorescence from amplification exceeded the background fluorescence values was set within the exponential growth region of the amplification curve. Real-time quantitative PCR data were analyzed using the comparative C_T_ method, as described by (Schmittgen and Livak, 2008). In each sample, we calculated the difference between the C_T_ values (ΔC_T_) for *Drd1a, Drd2* or *Slc1a1* and the housekeeping gene *Hprt*. Then, we calculated the difference in the ΔC_T_ values between the test and NTC samples and calculated 2^-ΔCT^. The ratio of these values was used to calculate the relative expression of different genes in samples from WT and EAAC1^−/−^ mice (**Fig. 6**).

**Figure 6.**
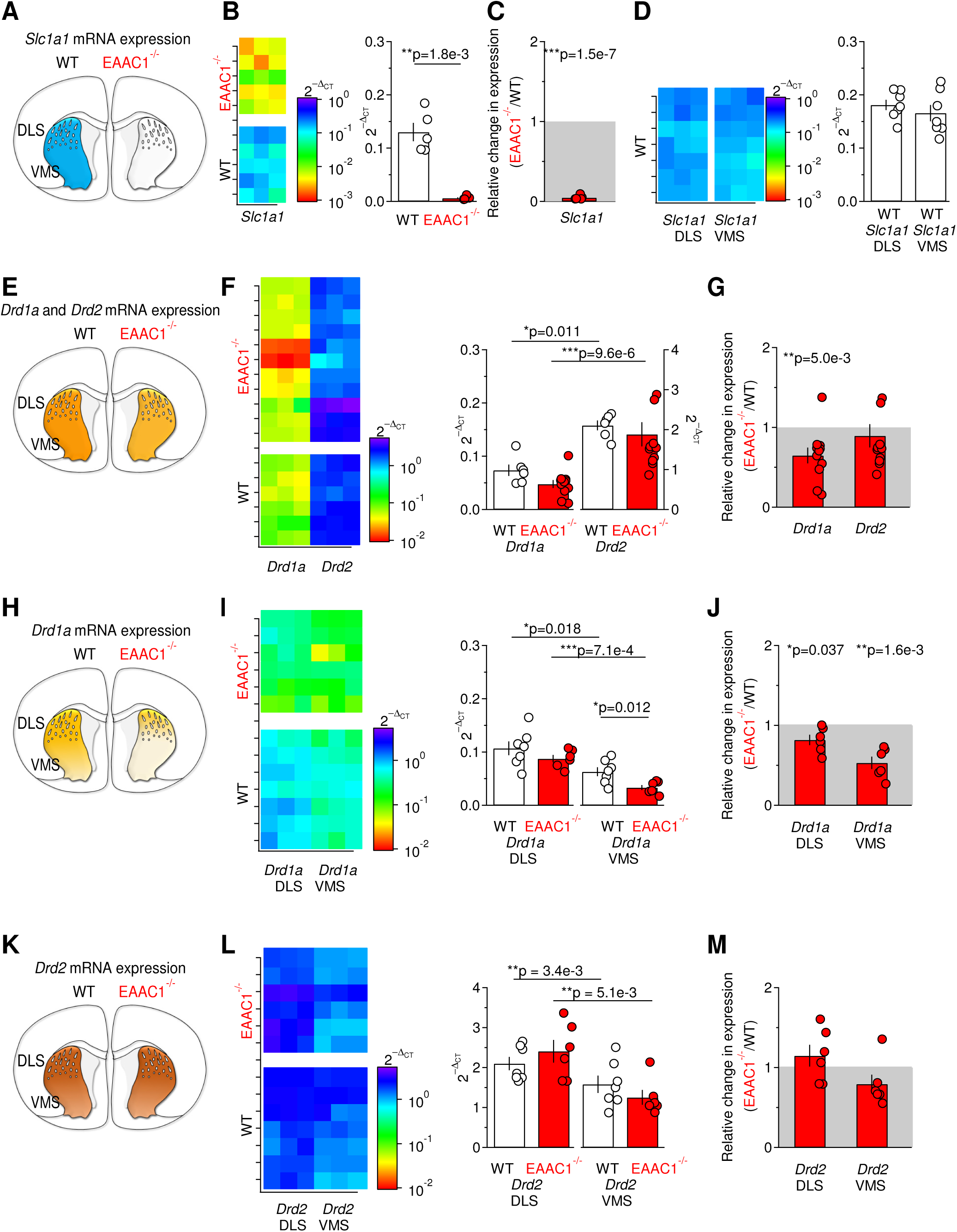
The mRNA levels of D1R are reduced in the DLS and VMS of EAAC1^−/−^ mice. **(A)** Schematic representation of a coronal section of the mouse brain. The blue region defines the WT striatum, in which the *Slc1a1* gene encoding EAAC1 is abundantly expressed. **(B)** Left: Color map representation of the *Slc1a1* levels in striatal samples from WT (n=5) and EAAC1^−/−^ mice (n=5), measured in qRT-PCR experiments. Each sample was run in triplicates (x-axis). Right: Relative amount of the *Slc1a1* gene with respect to the *Hprt* housekeeping gene in WT and EAAC1^−/−^ mice, measured as 2^-ΔCT^ (**p=1.8e-3). **(C)** Fold change in *Slc1a1* expression in EAAC1^−/−^ versus WT mice (2^-ΔCT^). Negligible levels of *Slc1a1* (***p=1.5e-7) are detected in EAAC1^−/−^ mice. **(D)** Left: Same experiments as described in (B, left), performed on samples from the DLS or VMS of WT mice (n=7). Right: As in (B, right), for samples from the DLS and VMC. No significant difference was detected in the expression levels of *Slc1a1* between the WT DLS and VMS (p=0.42). **(E)** Schematic representation of a coronal section of the mouse brain in which the striatum, from which we extracted the mRNA, is highlighted in yellow. The lighter shade of yellow indicates lower *Drd1a* gene expression levels in EAAC1^−/−^ mice. **(F)** Left: color map representation of the *Drdla* and *Drd2* levels in WT (n=6) and EAAC1^−/−^ mice (n=10), measured in qRT-PCR experiments. Each sample was run in triplicates (x-axis). Right: relative amount of the *Drd1a* and *Drd2* gene with respect to the *Hprt* housekeeping gene in WT and EAAC1^−/−^ mice, measured as 2^-ΔCT^. Higher levels of *Drd2* are detected in WT (*p=0.011) and EAAC1^−/−^ mice (***p=9.6e-6). (**G**) Fold change in *Drd1a* and *Drd2* expression in EAAC1^−/−^ vs WT mice (2^-ΔCT^). Reduced levels of *Drd1a* (**p=5.0e-3), not *Drd2* (p=0.48), are detected in EAAC1^−/−^ mice. **(H-J)** As in (E-G), on samples from the DLS and VMS of WT (n=7) and EAAC1^−/−^ mice (n=6). Lower levels of *Drd1a* were detected in the VMS of WT (*p=0.018) and EAAC1^−/−^ mice (***p_7 le_4). In the VMS, there was a significant reduction in the level of *Drd1a* (*p=0.012). A significant WT-fold change inDrd1a was detected in the DLS (*p=0.037) and VMS (**p=1.6e-3) of EAAC1^−/−^ mice. **(K-M)** As in (E-GF), for *Drd2*. The level of *Drd2* was lower in the VMS compared to the DLS, in WT (n=7, **p=3.4e-3) and EAAC1^−/−^ mice (n=6, **p=5.1e-3). No significant WT-fold change was detected in the expression level of *Drd2* in the DLS and VMS of EAAC1^−/−^ mice.

## Fluorescence labeling and imaging

D1^Cre/+^:Ai9^Tg/0^, D1_tdTomato/+_, A2A^Cre/+^:Ai9^Tg/0^, D1^Cre/+^:Ai9^Tg/O^:EAAC1^−/−^, D1^tdTomato/+^:EAAC1^−/−^, A2A^Cre/+^:Ai9^Tg/0^:EAAC1^−/−^ mice (P21-37) were deeply anesthetized with an I/P injection of pentobarbital (4 mg/g (w/w); Cat# 54925-045-10, Med-Pharmex, Pomona, CA) and trans-cardially perfused with 20 ml phosphate-buffered saline (PBS) 0.1 M and 20 ml 4% paraformaldehyde in PBS (4% PFA/PBS) at 4°C. The dissected brains were post-fixed overnight at 4°C in 4% PFA/PBS and cryo-protected at 4°C in 30% sucrose/PBS. Coronal sections (40 μm thick) were prepared using a vibrating blade microtome (VT1200S, Leica Microsystems, Wetzlar, Germany). All sections were post-fixed for 20 min at 4°C in 4% PFA/PBS. Sections used for cell density measures were then washed in PBS and mounted onto microscope slides using ProLong Gold anti-fade mounting medium (Cat #P36934, Thermo Fisher Scientific, Waltham, MA) or DAPI Fluoromount-G (Cat #0100-20, Southern Biotech, Birmingham, AL). The sections were then blocked/permeabilized for 1 hour at room temperature (RT) in a solution of PBS containing horse serum 10%, bovine serum albumin (BSA) 3% and Triton X-100 0.5%. The primary antibody incubation was performed by incubating the sections overnight at 4°C in a solution of PBS containing horse serum 3%, BSA 1% and Triton X-100 0.3% and one or more of the following primary antibodies: polyclonal rabbit anti-D1- or D2 dopamine receptor (1:100, Cat# ADR-001 and ADR-002, respectively; Alomone Labs, Jerusalem, Israel). The secondary antibody incubation was performed for 2 hours at RT using Alexa Fluor 488 goat anti-rabbit IgG (Cat# A-11034; Thermo Fisher Scientific) diluted to 1:500 in 0.1% Triton X-100/PBS. The brain sections were mounted onto microscope slides using ProLong Gold antifade mountant (Cat# P36934, Thermo Fisher Scientific, Waltham, MA) or DAPI Fluoromount-G (Cat #0100-20, Southern Biotech, Birmingham, AL). Confocal images were acquired using a Zeiss LSM710 inverted microscope equipped with 405 nm diode, 488 nm Ar and 561 nm DPSS lasers. All images (1024x1024 pixels) were acquired using a 63X oil immersion objective (N.A.=1.4) as averages of 16 consecutive images.

The image analysis was performed using the software Fiji (http://fiji.se/). Cell density measures were obtained by counting the number of immuno-labelled D1R- and D2R-expressing medium spiny neurons (D1- and D2-MSNs, respectively) in matrix areas of the DLS and VMS. The proportion of D1- and D2-MSNs was calculated as the ratio of D1^Cre/+^:Ai9^Tg/0^ or A2A^Cre/+^:Ai9^Tg/0^ and the total number of cell nuclei labelled with DAPI (**Supplementary Fig. 7-2**). The n-values reported in the text refer to the number of mice used for these experiments. Data were collected from at least 3 sections from each mouse.

## Western blot

Western blot experiments were performed on protein extracts from the striatum of WT and EAAC1^−/−^ mice of either sex (P21-36). Membrane and cytoplasmic protein extracts were obtained using the Mem-PER Plus Membrane Protein Extraction Kit (Cat# 89842; Thermo Fisher Scientific, Waltham, MA) according to the manufacturer’s instructions, using a cocktail of protease and phosphatase inhibitors (10 μl/ml, Cat# 78441; Thermo Fisher Scientific, Waltham, MA). The membrane protein extracts were used to measure protein levels of D1R and D2R (**Fig. 7,10,11; Supplementary Fig. 7-1**), mGluRI (**Fig. 4K**), GLAST and GLT-1 (data not shown). The cytoplasmic protein extracts were used to measure protein levels of the Dopamine- And cAMP-Regulated Phospho Protein 32 kDa (DARPP-32) and of its phosphorylated version (pDARPP-32; **Fig. 7,10,11**). The protein concentration was determined using spectrophotometer measures. We loaded equal amounts of proteins from WT and EAAC1^−/−^ mice in acrylamide gels (50-100 μg proteins in 10-12% acrylamide gels). The proteins were blotted onto polyvinylidene difluoride (PVDF) membranes (Cat# P2563; Millipore Sigma, St. Louis, MO) using a semi-dry blotting approach. The membranes were then blocked with 5% non-fat milk in TBST (pH=7.6) and probed using a primary antibody solution in which milk was replaced by BSA (5% BSA in TBST; pH=7.6). We used the following primary antibodies: rabbit anti D1R and D2R (1:200, Cat# ADR-001 and ADR-002 respectively; Alomone Labs, Jerusalem, Israel); rabbit anti mGluR5/1a (1:500, Cat# 2032-mGluR5/1a; PhosphoSolutions, Aurora, CO); rabbit anti GLAST (1:1,000, Cat# 5684; Cell Signaling Technology, Danvers, MA); rabbit anti GLT-1 (1:1,000, Cat# 3838; Cell Signaling Technology, Danvers, MA); rabbit anti DARPP-32 (1:1,000, Cat# 2306; Cell Signaling Technology, Danvers, MA); rabbit antibodies against different pDARPP-32 isoforms (pDARPP-32^T34^, 1:500, Cat# 12438; Cell Signaling Technology, Danvers, MA), (pDARPP-32^T75^ and pDARPP-32^S97^, 1:1,000, Cat# 2301 and Cat# 3401 respectively; Cell Signaling Technology, Danvers, MA), (pDARPP-32^S130^, 1:500, Cat# p1025-137; PhosphoSolutions, Aurora, CO), mCherry (Cat# 600-401-P16 Rockland Antibodies & Assays, Limerick, PA) and β-actin (1:1,000, Cat# 4970, Cell Signaling Technology, Danvers, MA). The membranes were incubated with the primary antibodies overnight. The pDARPP-32^S130^ antibody was incubated at RT. All other antibodies were incubated at 4°C. The secondary antibody incubation (biotinylated horse anti-rabbit IgG, Cat# BA-1100; Vector Laboratories, Burlingame, CA) was performed for 1-2 hours at RT with 5% non-fat milk in TBST (pH=7.6). The following secondary antibody dilutions were used for different proteins (1:200 pDARPP- 32^T34^; 1:500 mCherry; 1:1,000 for D2R, GLAST, GLT-1, pDARPP-32^S130^; 1:2,000 mGluRI; 1:3,000 pDARPP- 32^T75^; 1:4,000 D1R; 1:5,000 pDARPP-32^S97^, β-actin). Pre-adsorption experiments were performed using the control antigen provided by the supplier of the primary antibodies (Alomone Labs, Jerusalem, Israel) according to the manufacturer’s instructions (1 μg/ μg antibody). We amplified the immuno-labeling reactions with the Vectastain ABC kit (1:1,000 for pDARPP-32^T34^ and 1:2,000 for all other proteins, Cat# PK-6100; Vector Laboratories, Burlingame, CA) and the Clarity Western ECL (Cat# 170-5060; Bio-Rad, Hercules, CA) as a substrate for the peroxidase enzyme. For semi-quantitative analysis, protein band images were collected as 16 bit-images using a digital chemiluminescence imaging system (ChemiDoc, Bio-Rad, Hercules, CA or c300, Azure Biosystems, Dublin, CA) at different exposures (0.5-200 s). Each image was converted to an 8-bit image for image analysis, which was performed using the software Fiji (http://fiji.se/). Only images collected at exposure times that did not lead to pixel saturation were included in the analysis. The intensity of each band was calculated as the mean gray value in a region of interest (ROI) surrounding each band of interest, in three images collected using different exposure times. All band intensity values were normalized for the band intensity of β-actin in the same sample.

**Figure 4.**
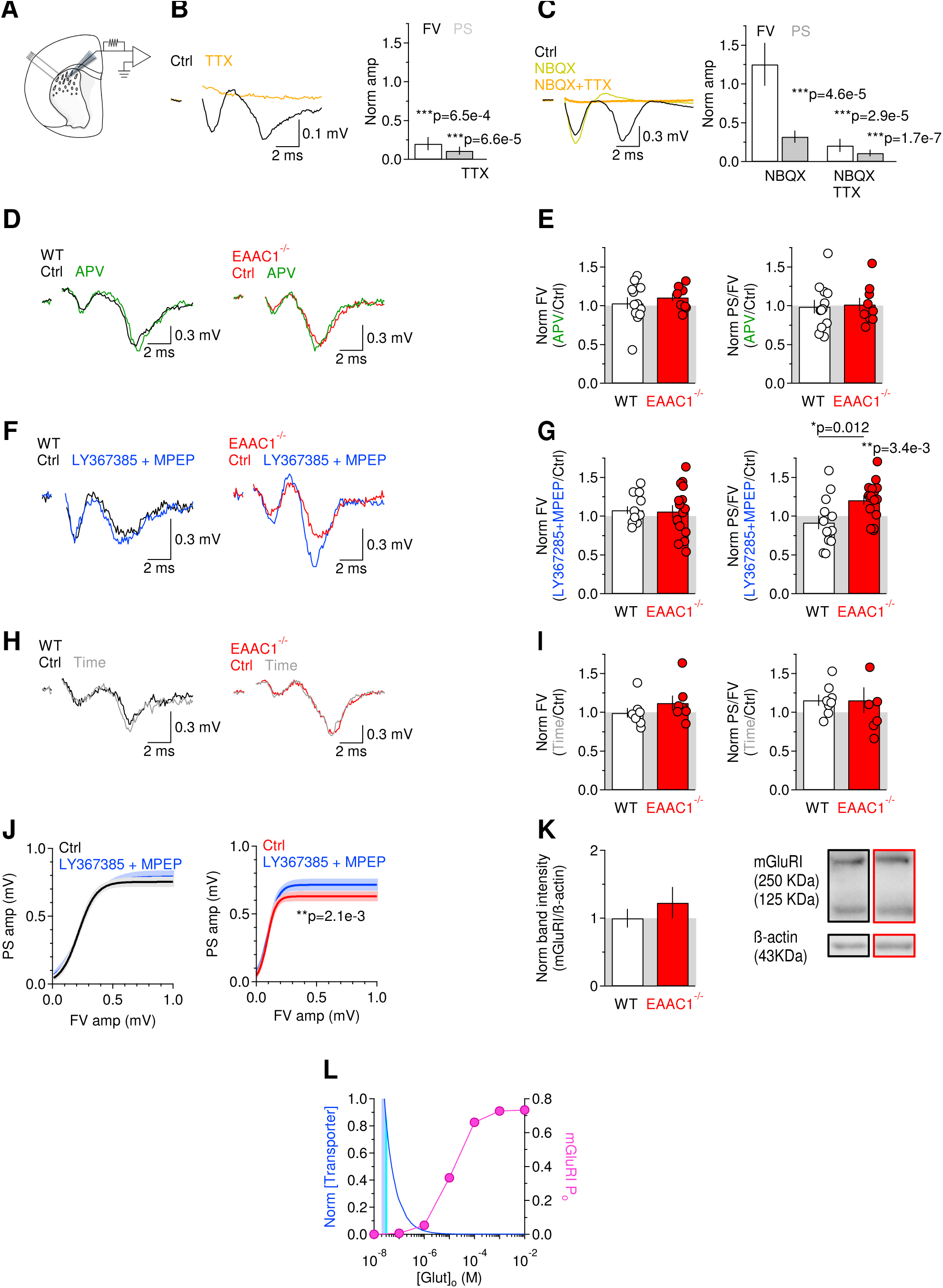
Excitatory field recordings in the DLS of EAAC1^−/−^ mice show increased sensitivity to mGluRI blockers. **(A)** Schematic illustration of the experimental design. The bipolar stimulating electrode and the extracellular field recording electrode were positioned in the DLS. **(B)** Left: extracellular recordings of FV and PS in the DLS of WT mice, in control conditions (black trace) and in the presence of the voltage-gated sodium channel blocker TTX (1 μM, orange trace). Right: Summary graph showing the effect of TTX on the FV amp (white bar (n=5), ***p=6.5e-4) and PS amplitude (grey bar (n=5), ***p=6.6e-5). **(C)** Left: extracellular recordings in the DLS of WT mice, in control conditions (black trace), in the presence of the GluA antagonist NBQX (10 μM, yellow trace) and in the additional presence of TTX (1 μM, orange trace). Right: summary graph showing that NBQX reduces the PS amplitude (grey bars (n=8), ***p=4.6e-5) without affecting the FV amplitude (white bars, p=0.39). The FV and PS are blocked in the presence of both NBQX and TTX (***p=2.9e-5 and ***p=1.7e-7, respectively). **(D)** Left: extracellular recordings in the DLS of WT mice, in control conditions (black trace) and in the presence of the GluN antagonist APV (50 μM, green trace). Right: extracellular recordings in the DLS of EAAC1^−/−^ mice, in control conditions (red trace) and in the presence of APV (50 μM, green trace). Each trace represents the average of 20 consecutive sweeps. **(E)** Left: Summary graph showing the effect of APV on the FV in WT (n=12) and EAAC1^−/−^ mice (n=9, WT vs. EAAC1^−/−^ p=0.41). Right: Effect of APV on the PS/FV ratio recorded in WT (n=12) and EAAC1^−/−^ mice (n=9, WT vs. EAAC1^−/−^ p=0.37). **(F)** As in (D), for recordings obtained in control conditions and in the presence of the mGluRI blockers LY367385 (50 μM) and MPEP (10 μM; blue traces). (**G**) Left: Summary graph showing the effect of LY367385 and MPEP on the FV in WT (n=13) and EAAC1^−/−^ mice (n=16, WT vs. EAAC1^−/−^ p=0.85). Right: Effect of LY367385 and MPEP on the PS/FV ratio recorded in WT (n=13) and EAAC1^−/−^ mice (n=16, WT vs. EAAC1^−/−^ *p=0.012). **(H)** As in (D), for recordings obtained in time-dependent control experiments. **(I)** Left: Summary graph showing the effect on the FV in WT (n=8) and EAAC1^−/−^ mice (n=7, WT vs. EAAC1^−/−^ p=0.28). Right: Effect on the PS/FV ratio recorded in WT (n=8) and EAAC1^−/−^ mice (n=7, WT vs. EAAC1^−/−^ p=1.00). **(J)** Input-output relationship between the PS and FV amplitudes evoked by stimulating excitatory afferents to the DLS. Left: mGluRI block does not alter the input-output curves in WT mice (n=13, p=0.57). Right: mGluRI block increases the PS amp over a range of stimulus intensities and FV amplitudes in EAAC1^−/−^ mice (n=16, **p=2.1e-3). **(K)** Western blot analysis showing similar levels of mGluRI expression in WT (n=6) and EAAC1^−/−^ mice (n=8; p=0.41). **(L)** We used a kinetic model of mGluRI (Marcaggi et al., 2009) to determine the open probability of these receptors at a range of extracellular glutamate concentrations (pink). The striatum maintains the extracellular glutamate concentration at ∼25 nM (Chiu and Jahr, 2017) in the presence of ∼140 μM glutamate transporters (Lehre and Danbolt, 1998). Reducing the concentration of glutamate transporters leads to increased extracellular glutamate concentrations (blue). EAAC1 accounts for less than 5-10% of all glutamate transporters. According to our model (see Methods), loss of EAAC1 can cause less than 10 nM increase in the ambient glutamate concentration (cyan). This change in the extracellular glutamate concentration can only cause a marginal increase in the mGluRI open probability.

**Figure 7.**
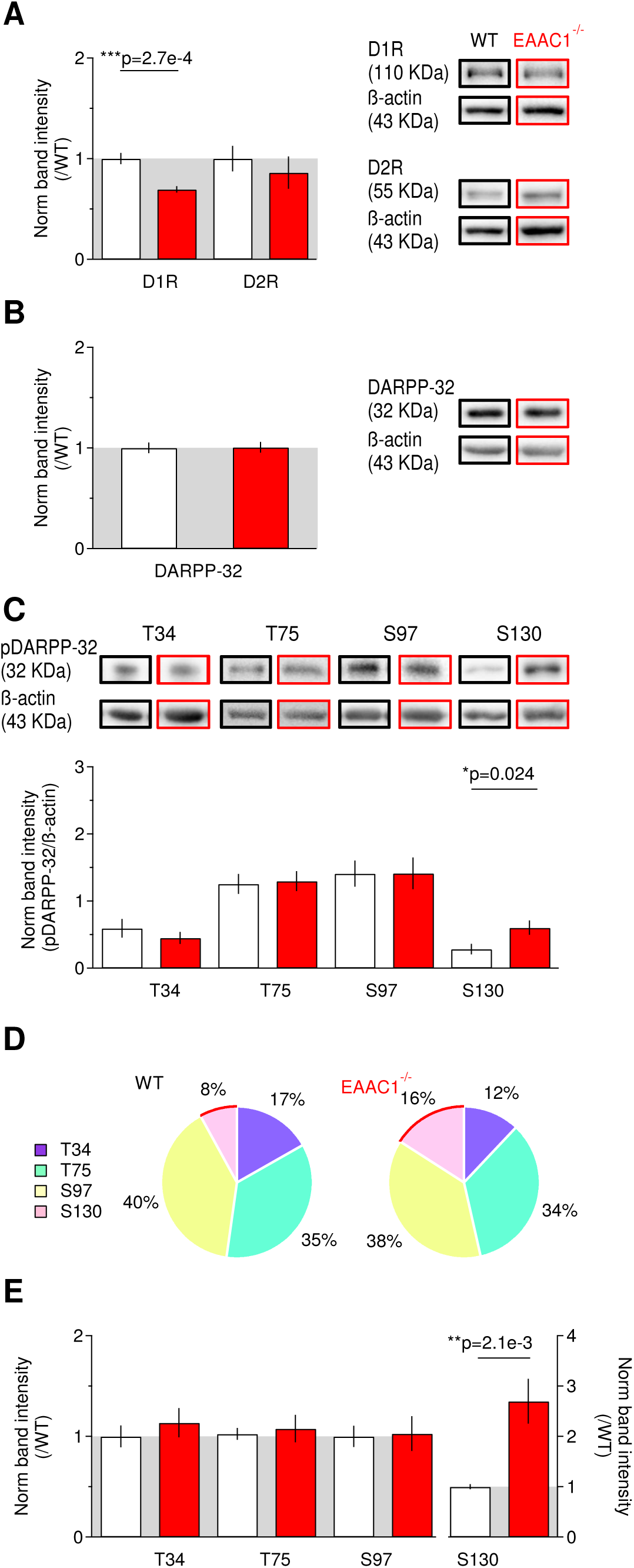
The reduced expression of D1R in EAAC1^−/−^ mice is associated with altered phosphorylation of DARPP-32. **(A)** Western blots for D1R, D2R and ß-actin in WT (n=ll) and EAAC1^−/−^ mice (n=8) show decreased levels of D1R (***p=2.7e-4), not D2R (p=0.51), in EAAC1^−/−^ mice. **(B)** Western blot analysis for DARPP-32, showing no significant difference in its expression between WT (n=15) and EAAC1^−/−^ mice (n=12, p=0.92). **(C)** Western blot analysis for pDARPP-32^T34^ (WT (n=10), EAAC1^−/−^ (n=10), p=0.41), pDARPP-32^175^ (WT (n=13), EAAC1^−/−^ (n=14), p=0.84), pDARPP-32^S97^ (WT (n=13), EAAC1^−/−^ (n=13), p=0.98), pDARPP-32^S130^ (WT (n=14), EAAC1^−/−^ (n=17), *p=0.024). **(D)** Pie chart representation of the phosphorylation distribution on the T34, T75, S97 and S130 sites of DARPP-32. The red curve highlights the S130 site, which shows increased phosphorylation in EAAC1^−/−^ mice. **(E)** As in (C), following data normalization by the average band intensity values measured in WT mice (pDARPP-32^T34^ WT (n=10), EAAC1^−/−^ (n=10), p=0.46; pDARPP-32^T75^ WT (n=13), EAAC1^−/−^ (n=14), p=0.72; pDARPP-32^S97^ WT (n=13), EAAC1^−/−^ (n=13), p=0.89; pDARPP-32^5130^ WT (n=14), EAAC1^−/−^ (n=15), **p=2.1e-3). Data in panels **?,?,?** represent the band intensity ratio between the target protein and β-actin in samples from WT versus EAAC1^7^' mice in the same blotting membrane.

**Figure 10.**
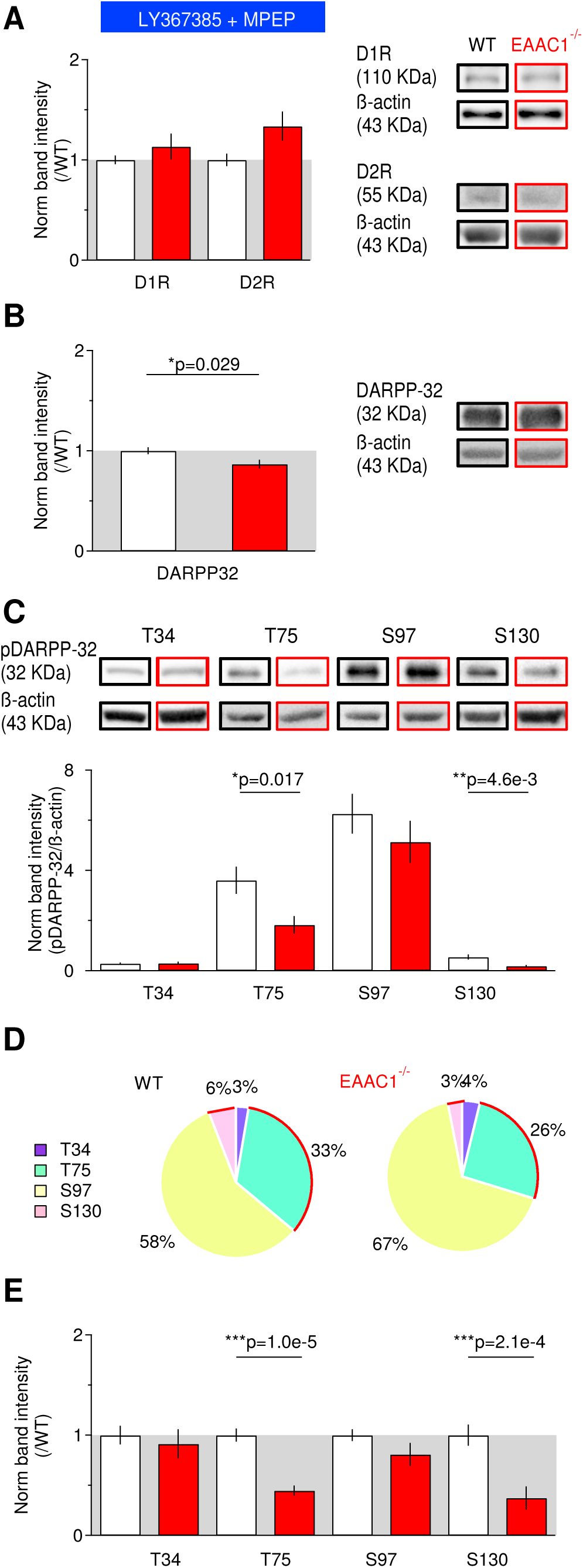
Blocking mGluRI activation rescues D1R expression and alters the phosphorylation pattern of DARPP-32. **(A)** Western blots for D1R, D2R and β-actin, in the presence of the mGluRI blockers LY367385 (50 μM) and MPEP (10 μM), show no significant difference in the expression of D1R and D2R (D1R: WT (n=9), EAAC1^−/−^ (n=9), p=0.35; D2R: WT (n=7), EAAC1^−/−^ (n=7), p=0.064). **(B)** mGluRI blockade induces a slight reduction in DARPP-32 expression between WT (n=9) and EAAC1^−/−^ mice (n=9, *p=0.029). (C) Western blot analysis for pDARPP-32^T34^ (WT (n=5), EAAC1^−/−^ (n=6), p=0.98), pDARPP-32^T75^ (WT (n=6), EAAC1^−/−^ (n=7), *p=0.017), pDARPP-32^S97^ (WT (n=9), EAAC1^−/−^ (n=9), p=0.34), pDARPP-32^S130^ (WT (n=12), EAAC1^−/−^ (n=9), **p=4.6e-3). **(D)** Pie chart representation of the phosphorylation distribution on the T34, T75, S97 and S130 sites of DARPP-32 in the presence of mGluRI blockers. The red curves highlight the T75 and S130 site, which show reduced phosphorylation in EAAC1^−/−^ mice. **(E)** As in (C), following data normalization by the average band intensity values measured in WT mice (WT (n=5), EAAC1^−/−^ (n=6), p=0.62), pDARPP-32^T75^ (WT (n=9), EAAC1^−/−^ (n=7), ***p=1.0e-5), pDARPP-32^S97^ (WT (n=9), EAAC1^−/−^ (n=9), p=0.16), pDARPP-32^S130^ (WT (n=12), EAAC1^−/−^ (n=9), ***p=2.1e-4). Data in panels **A,B,E** represent the band intensity ratio between the target protein and β-actin measured in samples from EAAC1^−/−^ mice and normalized by analogous measures in samples from WT mice blotted in the same membrane.

**Figure 11.**
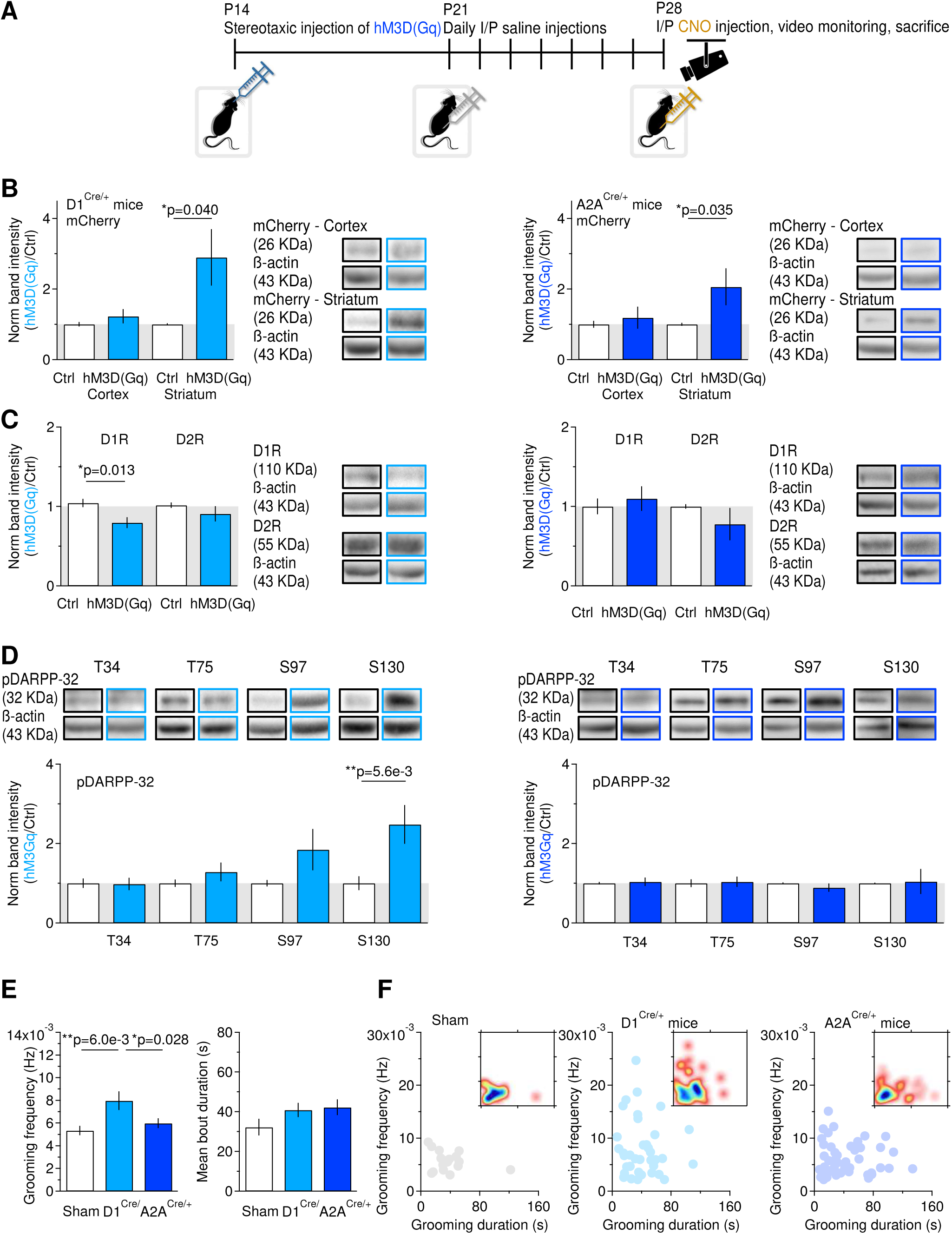
Cell-specific activation of Gq signaling pathway induces decreased D1R expression and increased pDARPP-32^S130^ phosphorylation. **(A)** Timeline of the experimental design. At P14-16, mice received a unilateral stereotaxic injection of hM3D(Gq). After one week, they started receiving daily I/P saline injections. At P28-30, they received I/P injections of CNO (5 mg/Kg). One hour after the CNO injections, they were video-monitored to examine their grooming behavior. Two hours after the CNO injections, they were sacrificed. Proteins for Western blot analysis were extracted from the control and injected striatum and from the adjacent cortices. **(B)** Left: mCherry expression in D1^Cre/+^ mice. A significant increase in mCherry expression was detected only in the striatum of D1^Cre/+^ mice (n=10, *p=0.040). Right: mCherry expression in A2A^Cre/+^ mice. A significant increase in mCherry expression was detected only in the striatum of A2A^Cre/+^ mice (n=23, *p=0.035). **(C)** Left: D1R and D2R expression in D1^Cre/+^ mice (D1R: n=9, *p=0.013; D2R: n=8, p=0.35). The expression of D1R is significantly reduced in the injected striatum. Right: D1R and D2R expression in A2A^Cre/+^ mice (D1R: n=9, p=0.63; D2R: n=5, p=0.34). The expression of D1R and D2R is similar in the injected and non-injected striatum. **(D)** Left: hM3D(Gq) injection in D1^Cre/+^ mice leads to increased pDARPP-32^S130^ (pDARPP-32^T34^ n=10, p=0.92; pDARPP-32^T75^ n=11, p=0.18; pDARPP-32^S97^ n=11, p=0.13, pDARPP-32^S130^ n=11, **p=5.6e-3). Right: hM3D(Gq) injection in A2A^Cre/+^ mice leads to no change in pDARPP-32 (pDARPP-32^T34^ n=9, p=0.75; pDARPP-32^T75^ n=8, p=0.84; pDARPP-32^S97^ n=8, p=0.31, pDARPP-32^S130^ n=6, p=0.89). (E) Summary of the frequency (Frequency: Sham (n=25), D1^Cre/+^ (n=39), Sham vs D1^Cre/+^ **p=6.0e-3, A2A^Cre/+^ (n=49), Sham vs A2A^Cre/+^ p=0.28) and duration of grooming episodes in Sham and D1^Cre/+^ mice injected with hM3D(Gq) (Duration: Sham (n=34), D1^Cre/+^ (n=44), Sham vs D1^Cre/+^ p=0.12, A2A^Cre/+^ (n=58), Sham vs A2A^Cre/+^ p=0.08). **(F)** Relationship between the frequency and duration of grooming episodes in Sham (left), D1^Cre/+^ (middle) and A2A^Cre/+^ mice (right). The inset represents a density plot of the data, with blue areas corresponding to the duration and frequency of the most commonly observed grooming episodes.

## Acute slice preparation, electrophysiology experiments and data analysis

Acute coronal slices of the mouse striatum were obtained from WT and EAAC1^−/−^ mice of either sex (P16-24), deeply anesthetized with isoflurane and decapitated in accordance with SUNY Albany Animal Care and Use Committee guidelines. The brain was rapidly removed and placed in ice-cold slicing solution bubbled with 95% O_2_-5% CO_2_, containing (in mM): 119 NaCl, 2.5 KCl, 0.5 CaCl_2_, 1.3 MgSO_4_H_2_O, 4 MgCl_2_, 26.2 NaHCO_3_, 1 NaH_2_PO_4_, and 22 glucose; 320 mOsm; pH 7.4. The slices (250 μm thick) were prepared using a vibrating blade microtome (VT1200S, Leica Microsystems, Wetzlar, Germany). Once prepared, the slices were stored in this solution in a submersion chamber at 36°C for 30 min and at RT for at least 30 min and up to 5 hours. Unless otherwise stated, the recording solution contained (in mM): 119 NaCl, 2.5 KCl, 1.2 CaCl_2_, 1 MgCl_2_, 26.2 NaHCO_3_, 1 NaH_2_PO_4_, 22 glucose; 300 mOsm; pH 7.4. This solution was also used to fill the glass capillaries used to obtain extracellular field recordings. We identified the DLS under bright light illumination using an upright fixed-stage microscope (BX51 WI, Olympus Corporation, Tokyo, Japan). Stimulating and recording electrodes were both placed in the DLS, ∼100 μm away from each other. Post-synaptic responses were evoked by delivering constant voltage electrical pulses (50 μs) through a stimulating bipolar stainless steel electrode (Cat# MX21AES(JD3); Frederick Haer Company, Bowdoin, ME). The resistance of the recording electrode was ∼1.5 MΩ and was monitored throughout the experiments. Data were discarded if the resistance changed more than 20% during the course of the experiment. Picrotoxin (100 μM) was added to the recording solution to block GABA_A_ receptors. All recordings were obtained using a Multiclamp 700B amplifier and a 10 KHz low-pass filter (Molecular Devices, Sunnyvale, CA). All traces were digitized at 10 KHz and analyzed off-line with custom-made software (A.S.) written in IgorPro 6.36 (Wavemetrics, Lake Oswego, OR). Tetrodotoxin (TTX) was purchased from Alomone Labs (Jerusalem, Israel). 2,3-Dioxo-6-nitro-1,2,3,4-tetrahydrobenzo[f]quinoxaline-7-sulfonamide disodium salt (NBQX), DL-2-Amino-5- phosphonopentanoic acid (APV), (S)-(+)-α-Amino-4-carboxy-2-methylbenzeneaceticacid (LY367385) and 2-Methyl-6-(phenylethynyl)pyridine hydrochloride (MPEP) were purchased from Tocris Bioscience (Bristol, UK) and Hello Bio (Princeton, NJ). All other chemicals were purchased from Millipore Sigma (St. Louis, MO). All experiments were performed at RT.

## Stereotaxic intracranial injections

Male and female D1^Cre/+^and WT mice (P14-16) were anesthetized with isoflurane (induction: 5% in 100% 0_2_ at 1-2 l/min; maintenance: 3% in 100% O_2_ at 1-2 l/min) and placed in the stereotaxic frame of a motorized drill and injection robot (Neurostar, Tubingen, Germany). After making a skin incision and thinning the skull under aseptic conditions, we injected 100 nl of the AAV construct AAV-hSyn-DIO-hM3D(Gq)-mCherry (University of North Carolina, Chapel Hill, NC) unilaterally in either the left or right DLS using a Hamilton syringe at a rate of 100 nl/min. The non-injected striatum was used as an internal control. The injection coordinates from lambda were AP: -2.5 mm, ML: ±2.0 mm, DV: +5.0 mm. After the stereotaxic injections, the mice were returned to their home cage for 7 days. Then, they received daily I/P injections of NaCl 0.9% (10 μl/g, v/w) for 7 days. Two weeks after the stereotaxic surgery, mice received a single I/P injection of CNO (5 mg/Kg in NaCl 0.9%) (Enzo Life Sciences, Farmingdale, NY). One hour after the CNO injection, we acquired videos to monitor the grooming behavior of the mice. Two hours after the CNO injections, we euthanized the mice to isolate the left and right striatum and cortex to be used for Western blot analysis.

## Computer modeling

### Kinetic model of mGluRI

We used ChanneLab (Synaptosoft) to estimate the mGluRI open probability using a kinetic scheme of mGluRI activation (Marcaggi et al., 2009). An analytic approach was used to determine the relationship between glutamate transporter concentration and ambient glutamate concentration (**Fig. 4L**). Briefly, under steady-state conditions, the relationship between glutamate transporter and extracellular glutamate concentration can be described by a modified version of the Michaelis-Menten equation (Sun et al., 2014), as follows:

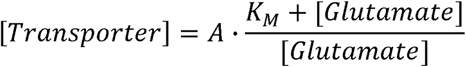

In this equation, *K_M_*=27 μM (Sun et al., 2014). The proportionality constant A can be calculated by setting [Glutamate]=25 nM (the experimentally measured ambient glutamate concentration in the striatum (Chiu and Jahr, 2017)) and [Transporter]=140 μM (the estimated concentration of glutamate transporters in the brain (Lehre and Danbolt, 1998)). The data shown in **Fig. 4L** (blue) were normalized to obtain the normalized transporter concentration (y-axis) at different steady state ambient glutamate concentrations (x-axis). The analysis of this relationship was performed using custom software written in IgorPro (Wavemetrics, Lake Oswego, OR; A.S.).

### 3D Monte Carlo model of pDARPP-32 phosphorylation

We used Blender (2.75) to create a simplified 3D mesh geometry which we used as an *in silico* representation of an excitatory post-synaptic terminal (i.e. spine head). This 3D geometry was shaped as a 1 μm^3^ volume sphere of radius *r*=0.62 μm, with reflective properties for any diffusing molecule (i.e. a diffusing molecule contacting the sphere bounces back without disappearing from the simulation environment). This geometry was exported into Model Description Language (MDL) and was used in the MCell (3.4) 3D Monte Carlo simulation environment. At the beginning of each simulation, we released DARPP-32 and other protein kinases and phosphatases in the sphere, at the concentration specified in **Table 6.** We let the system equilibrate for 150 s to allow all molecules to diffuse evenly and equilibrate throughout the entire simulation environment (i.e. the sphere). After this equilibration time, we released Ca^2+^ (1 μM-1 mM) from the center of the sphere using either single pulses or trains of 10 pulses (0.1-10 Hz) as described in **Fig. 8.** The diffusion coefficient of Ca^2+^ was set to *D\**=2.2e-6 cm^2^/s (Allbritton et al., 1992). The diffusion coefficient of all other molecules was set to *D\**=5e-8 cm^2^/s (Li et al., 2015). The initial conditions (**Table 6**) and reactions (**Table 7**) were modeled according to the kinetic schemes reported by (Fernandez et al., 2006), to which we added new reactions for pDARPP-32^S97^. Briefly, DARPP-32 is phosphorylated in position T34 by the cAMP-dependent protein kinase A (PKA; pDARPP- 32^T34^), in position T75 by the cyclin-dependent kinase 5 (CdK5; pDARPP-32^T75^), in position S97 by casein kinase 2 (CK2; pDARPP-32^S97^) and in position S130 by casein kinase 1 (CK1; pDARPP-32^S130^). The phosphorylation of each position was independent of the phosphorylation state of other positions, as specified by the available kinetic parameters. Each simulation was run 100 times. Each run lasted 500 s and consisted of 5,000 iterations with a time step *Δt*=0.1 s. The concentration of each molecule in the sphere was monitored at every *Δt*. We used custom-made scripts written in Python 3.5 (A.S.) to execute each run and average the results obtained over multiple runs, which are shown in **Fig. 8.** In this figure, pDARPP-32^S130^ represents the sum of all pDARPP-32 states that have one phosphate group attached to position S130, regardless of the phosphorylation state of other sites.

**Table 6.** Initial conditions used in the models

| Molecule name | Number of molecules | Concentration (μM) |
| --- | --- | --- |
| DARPP-32 | 3,000 | 4.98 |
| Cdk5 | 120 | 0.2 |
| CK1 | 100 | 0.166 |
| CK2 | 100 | 0.166 |
| PDE | 1,204 | 2 |
| PP2A | 120 | 0.2 |
| PP2B (inactive) | 200 | 0.332 |
| PP2C | 80 | 0.133 |
| R2C2 | 4,000 | 6.64 |
| Ca<sup>2+</sup> | 60 | 0.1 |

**Table 7.** Reaction parameters used in the models

| Reaction | $k_{on} (M^{-1} s^{-1})$ | $k_{off} (s^{-1})$ | $\alpha (s^{-1})$ |
| --- | --- | --- | --- |
| DARPP-32 + Cdk5 → DARPP-32-Cdk5 | 5.6e+06 | 12 |  |
| DARPP-32-Cdk5 → DARPP-32<sup>T75</sup> + Cdk5 |  | 3 |  |
| DARPP-32 + CK1 → DARPP-32-CK1 | 4.4e+06 | 12 |  |
| DARPP-32-CK1 → DARPP-32<sup>S130</sup> + CK1 |  | 3 |  |
| DARPP-32 + PKA → DARPP-32-PKA | 5.6e+06 | 10.8 |  |
| DARPP-32-PKA → DARPP-32<sup>T34</sup> + PKA |  | 2.7 |  |
| DARPP-32<sup>T34</sup> + Cdk5 → DARPP-32<sup>T34</sup>-Cdk5 | 5.6e+06 | 12 |  |
| DARPP-32<sup>T34</sup>-Cdk5 → DARPP-32<sup>T34,T75</sup> + Cdk5 |  | 3 |  |
| DARPP-32<sup>T34</sup> + CK1 → DARPP-32<sup>T34</sup>-CK1 | 4.4e+06 | 12 |  |
| DARPP-32<sup>T34</sup>-CK1 → DARPP-32<sup>T34,S130</sup> + CK1 |  | 3 |  |
| DARPP-32<sup>T34</sup> + PP2B → DARPP-32<sup>T34</sup>-PP2B | 1e+08 | 16 |  |
| DARPP-32<sup>T34</sup>-PP2B → DARPP-32 + PP2B |  | 4 |  |
| DARPP-32<sup>T75</sup> + CK1 → DARPP-32<sup>T75</sup>-CK1 | 4.4e+06 | 12 |  |
| DARPP-32<sup>T75</sup>-CK1 → DARPP-32<sup>T75,S130</sup> + CK1 |  | 3 |  |
| DARPP-32<sup>T75</sup> + PKA → DARPP-32<sup>T75</sup>-PKA | 5.6e+06 | 10.8 |  |
| DARPP-32<sup>T75</sup>-PKA → DARPP-32<sup>T34,T75</sup> + PKA |  | 0 |  |
| DARPP-32<sup>T75</sup> + PP2A → DARPP-32<sup>T75</sup>-PP2A | 3.8e+06 | 24 |  |
| DARPP-32<sup>T75</sup>-PP2A → DARPP-32 + PP2A |  | 10 |  |
| DARPP-32<sup>T75</sup> + PP2AP → DARPP-32<sup>T75</sup>-PP2AP | 1.7e+07 | 24 |  |
| DARPP-32<sup>T75</sup>-PP2AP → DARPP-32<sup>T75</sup> + PP2AP |  | 40 |  |
| DARPP-32<sup>S130</sup> + Cdk5 → DARPP-32<sup>S130</sup>-Cdk5 | 5.6e+06 | 12 |  |
| DARPP-32<sup>S130</sup>-Cdk5 → DARPP-32<sup>T75,S130</sup> + Cdk5 |  | 3 |  |
| DARPP-32<sup>S130</sup> + PKA → DARPP-32<sup>S130</sup>-PKA | 5.6e+06 | 10.8 |  |
| DARPP-32<sup>S130</sup>-PKA → DARPP-32<sup>T34,S130</sup> + PKA |  | 2.7 |  |
| DARPP-32<sup>S130</sup> + PP2C → DARPP-32<sup>S130</sup>-PP2C | 7.5e+06 | 12 |  |
| DARPP-32<sup>S130</sup>-PP2C → DARPP-32 + PP2C |  | 3 |  |
| DARPP-32<sup>T34,T75</sup> + CK1 → DARPP-32<sup>T34,T75</sup>-CK1 | 4.4e+06 | 12 |  |
| DARPP-32<sup>T34,T75</sup>-CK1 → DARPP-32<sup>T34,T75,S130</sup> + CK1 |  | 3 |  |
| DARPP-32<sup>T34,T75</sup> + PP2A → DARPP-32<sup>T34,T75</sup>-PP2A | 3.8e+06 | 24 |  |
| DARPP-32<sup>T34,T75</sup>-PP2A → DARPP-32<sup>T34</sup> + PP2A |  | 10 |  |
| DARPP-32<sup>T34,T75</sup> + PP2AP -> DARPP-32<sup>T34,T75</sup>-PP2AP | 1.7e+07 | 40 |  |
| DARPP-32<sup>T34,T75</sup>-PP2AP -> DARPP-32<sup>T34</sup> + PP2AP |  | 24 |  |
| DARPP-32<sup>T34,T75</sup> + PP2B -> DARPP-32<sup>T34,T75</sup>-PP2B | 1e+08 | 1, 600 |  |
| DARPP-32<sup>T34,T75</sup>-PP2B -> DARPP-32<sup>T75</sup> + PP2B |  | 4 |  |
| DARPP-32<sup>T34,S130</sup> + CdK5 -> DARPP-32<sup>T34,S130</sup>-CdK5 | 5.6e+06 | 12 |  |
| DARPP-32<sup>T34,S130</sup>-CdK5 -> DARPP-32<sup>T34,T75,S130</sup> + CdK5 |  | 3 |  |
| DARPP-32<sup>T34,S130</sup> + PP2B -> DARPP-32<sup>T34,S130</sup>-PP2B | 7.5e+04 | 0.12 |  |
| DARPP-32<sup>T34,S130</sup>-PP2B -> DARPP-32<sup>S130</sup> + PP2B |  | 0.03 |  |
| DARPP-32<sup>T34,S130</sup> + PP2C -> DARPP-32<sup>T34,S130</sup>-PP2C | 7.5e+06 | 12 |  |
| DARPP-32<sup>T34,S130</sup>-PP2C -> DARPP-32<sup>T34</sup> + PP2C |  | 3 |  |
| DARPP-32<sup>T75,S130</sup> + PKA -> DARPP-32<sup>T75,S130</sup>-PKA | 5.6e+06 | 10.8 |  |
| DARPP-32<sup>T75,S130</sup>-PKA -> DARPP-32<sup>T34,T75,S130</sup> + PKA |  | 0 |  |
| DARPP-32<sup>T75,S130</sup> + PP2A -> DARPP-32<sup>T75,S130</sup>-PP2A | 3.8e+06 | 24 |  |
| DARPP-32<sup>T75,S130</sup>-PP2A -> DARPP-32<sup>S130</sup> + PP2A |  | 10 |  |
| DARPP-32<sup>T75,S130</sup> + PP2AP -> DARPP-32<sup>T75,S130</sup>-PP2AP | 1.7e+07 | 40 |  |
| DARPP-32<sup>T75,S130</sup>-PP2AP -> DARPP-32<sup>S130</sup> + PP2AP |  | 24 |  |
| DARPP-32<sup>T75,S130</sup> + PP2C -> DARPP-32<sup>T75,S130</sup>-PP2C | 7.5e+06 | 12 |  |
| DARPP-32<sup>T75,S130</sup>-PP2C -> DARPP-32<sup>T75</sup> + PP2C |  | 3 |  |
| DARPP-32<sup>T34,T75,S130</sup> + PP2A -> DARPP-32<sup>T34,T75,S130</sup>-PP2A | 3.8e+06 | 24 |  |
| DARPP-32<sup>T34,T75,S130</sup>-PP2A -> DARPP-32<sup>T34,S130</sup> + PP2A |  | 10 |  |
| DARPP-32<sup>T34,T75,S130</sup> + PP2AP -> DARPP-32<sup>T34,T75,S130</sup>-PP2AP | 1.7e+07 | 40 |  |
| DARPP-32<sup>T34,T75,S130</sup>-PP2AP -> DARPP-32<sup>T34,S130</sup> + PP2AP |  | 24 |  |
| DARPP-32<sup>T34,T75,S130</sup> + PP2B -> DARPP-32<sup>T34,T75,S130</sup>-PP2B | 7.5e+04 | 120 |  |
| DARPP-32<sup>T34,T75,S130</sup>-PP2B -> DARPP-32<sup>T75,S130</sup> + PP2B |  | 0.03 |  |
| DARPP-32<sup>T34,T75</sup> + PP2C -> DARPP-32<sup>T34,T75,S130</sup>-PP2C | 7.5e+06 | 3 |  |
| DARPP-32<sup>T34,T75,S130</sup>-PP2C -> DARPP-32<sup>T34,T75,S130</sup> + PP2C |  | 12 |  |
| CK1P + PP2B -> CK1P-PP2B | 3e+08 | 24 |  |
| CK1P-PP2B -> CK1 + PP2B |  | 6 |  |
| CK1 -> CK1P |  |  | 1 |
| PDE + PKA -> PDE-PKA | 6e+06 | 36 |  |
| PDE-PKA -> PDEP + PKA |  | 9 |  |
| PDEP -> PDE |  |  | 0.1 |
| PP2A + PKA -> PP2A-PKA | 1e+08 | 16 |  |
| PP2A-PKA -> PP2AP + PKA |  | 4 |  |
| PP2AP -> PP2A |  |  | 5 |
| PP2B(inactive) + Ca<sup>2+</sup> -> PP2B(inactive)-Ca<sup>2+</sup> | 1e+15 | 1 |  |
| PP2B(inactive)-Ca<sup>2+</sup> + Ca<sup>2+</sup> -> PP2B | 3e+15 | 1 |  |
| R2C2 + cAMP -> cAMP-R2C2 | 5.4e+07 | 33 |  |
| cAMP-R2C2 + cAMP -> cAMP2-R2C2 | 5.4e+07 | 33 |  |
| cAMP2-R2C2 + cAMP -> cAMP3-R2C2 | 7.5e+07 | 110 |  |
| cAMP3-R2C2 + cAMP -> cAMP4-R2C2 | 7.5e+07 | 32.5 |  |
| cAMP4-R2C + PKA -> cAMP4-R2C2 | 1.8e+07 | 60 |  |
| cAMP4-R2 + PKA -> cAMP4-R2C | 1.8e+07 | 60 |  |
| cAMP + PDE -> cAMP-PDE | 2.52e+06 | 40 |  |
| cAMP-PDE -> AMP + PDE |  | 10 |  |
| cAMP + PDEP -> cAMP-PDEP | 5.04e+06 | 80 |  |
| cAMP-PDEP -> AMP + PDEP |  | 20 |  |
| Ca<sup>2+</sup> -> NULL |  |  | 0.6 |
| Ca<sup>2+</sup> + CK2 -> CK2-Ca<sup>2+</sup> | 1e+15 | 3e+15 |  |
| CK2-Ca<sup>2+</sup> + DARPP-32 -> DARPP-32-CK2-Ca<sup>2+</sup> | 4.4e5 | 12 |  |
| DARPP-32-CK2-Ca<sup>2+</sup> -> DARPP-32<sup>S97</sup> + CK2-Ca<sup>2+</sup> |  | 3 |  |
| DARPP-32<sup>S97</sup> + PP2A -> DARPP-32<sup>S97</sup>-PP2A | 3.8e+06 | 24 |  |
| DARPP-32<sup>S97</sup>-PP2A -> DARPP-32 + PP2A |  | 10 |  |
| DARPP-32<sup>T34,S97</sup> + PP2A -> DARPP-32<sup>T34,S97</sup>-PP2A | 3.8e+06 | 24 |  |
| DARPP-32<sup>T34,S97</sup>-PP2A -> DARPP-32<sup>T34</sup> + PP2A |  | 10 |  |
| DARPP-32<sup>T34,S97</sup>-PP2A -> DARPP-32<sup>S97</sup> + PP2A |  | 10 |  |
| DARPP-32<sup>T75,S97</sup> + PP2A -> DARPP-32<sup>T75,S97</sup>-PP2A | 3.8e+06 | 24 |  |
| DARPP-32<sup>T75,S97</sup>-PP2A -> DARPP-32<sup>T75</sup> + PP2A |  | 10 |  |
| DARPP-32<sup>T75,S97</sup>-PP2A -> DARPP-32<sup>S97</sup> + PP2A |  | 10 |  |
| DARPP-32<sup>S97,S130</sup> + PP2A -> DARPP-32<sup>S97,S130</sup>-PP2A | 3.8e+06 | 24 |  |
| DARPP-32<sup>S97,S130</sup>-PP2A -> DARPP-32<sup>S130</sup> + PP2A |  | 10 |  |
| DARPP-32<sup>S97,S130</sup>-PP2A -> DARPP-32<sup>S97</sup> + PP2A |  | 10 |  |
| DARPP-32<sup>T34, T75, S97</sup> + PP2A -> DARPP-32<sup>T34, T75, S97</sup>-PP2A | 3.8e+06 | 24 |  |
| DARPP-32<sup>T34, T75, S97</sup>-PP2A -> DARPP-32<sup>T75, S97</sup> + PP2A |  | 10 |  |
| DARPP-32<sup>T34, T75, S97</sup>-PP2A -> DARPP-32<sup>T34, S97</sup> + PP2A |  | 10 |  |
| DARPP-32<sup>T34, T75, S97</sup>-PP2A -> DARPP-32<sup>T34, T75</sup> + PP2A |  | 10 |  |
| DARPP-32<sup>T75, S97, S130</sup> + PP2A -> DARPP-32<sup>T75, S97, S130</sup>-PP2A | 3.8e+06 | 24 |  |
| DARPP-32<sup>T75, S97, S130</sup>-PP2A -> DARPP-32<sup>S97, S130</sup> + PP2A |  | 10 |  |
| DARPP-32<sup>T75, S97, S130</sup>-PP2A -> DARPP-32<sup>T75, S130</sup> + PP2A |  | 10 |  |
| DARPP-32<sup>T75, S97, S130</sup>-PP2A -> DARPP-32<sup>T75, S97</sup> + PP2A |  | 10 |  |
| DARPP-32<sup>T34, S97, S130</sup> + PP2A -> DARPP-32<sup>T34, S97, S130</sup>-PP2A | 3.8e+06 | 24 |  |
| DARPP-32<sup>T34, S97, S130</sup>-PP2A -> DARPP-32<sup>S97, S130</sup> + PP2A |  | 10 |  |
| DARPP-32<sup>T34, S97, S130</sup>-PP2A -> DARPP-32<sup>T34, S130</sup> + PP2A |  | 10 |  |
| DARPP-32<sup>T34, S97, S130</sup>-PP2A -> DARPP-32<sup>T34, S97</sup> + PP2A |  | 10 |  |
| DARPP-32<sup>T34, T75, S130</sup> + PP2A -> DARPP-32<sup>T34, T75, S130</sup>-PP2A | 3.8e+06 | 24 |  |
| DARPP-32<sup>T34, T75, S130</sup>-PP2A -> DARPP-32<sup>T75, S130</sup> + PP2A |  | 10 |  |
| DARPP-32<sup>T34, T75, S130</sup>-PP2A -> DARPP-32<sup>T34, S130</sup> + PP2A |  | 10 |  |
| DARPP-32<sup>T34, T75, S130</sup>-PP2A -> DARPP-32<sup>T34, T75</sup> + PP2A |  | 10 |  |
| DARPP-32<sup>T34, T75, S97, S130</sup> + PP2A -> DARPP-32<sup>T34, T75, S97, S130</sup>-PP2A | 3.8e+06 | 24 |  |
| DARPP-32<sup>T34, T75, S97, S130</sup>-PP2A -> DARPP-32<sup>T75, S97, S130</sup> + PP2A |  | 10 |  |
| DARPP-32<sup>T34, T75, S97, S130</sup>-PP2A -> DARPP-32<sup>T34, S97, S130</sup> + PP2A |  | 10 |  |
| DARPP-32<sup>T34, T75, S97, S130</sup>-PP2A -> DARPP-32<sup>T34, T75, S130</sup> + PP2A |  | 10 |  |
| DARPP-32<sup>T34, T75, S97, S130</sup>-PP2A -> DARPP-32<sup>T34, T75</sup> + PP2A |  | 10 |  |
| DARPP-32<sup>S97</sup> + CdK5 -> DARPP-32<sup>S97</sup>-CdK5 | 5.6e+06 | 12 |  |
| DARPP-32<sup>S97</sup>-CdK5 -> DARPP-32<sup>T75, S97</sup> + CdK5 |  | 3 |  |
| DARPP-32<sup>T34, S97</sup> + CdK5 -> DARPP-32<sup>T34, S97</sup>-CdK5 | 5.6e+06 | 12 |  |
| DARPP-32<sup>T34, S97</sup>-CdK5 -> DARPP-32<sup>T34, T75, S97</sup> + CdK5 |  | 3 |  |
| DARPP-32<sup>S97, S130</sup> + CdK5 -> DARPP-32<sup>S97, S130</sup>-CdK5 | 5.6e+06 | 12 |  |
| DARPP-32<sup>S97, S130</sup>-CdK5 -> DARPP-32<sup>T75, S97, S130</sup> + CdK5 |  | 3 |  |
| DARPP-32<sup>T34, S97, S130</sup> + CdK5 -> DARPP-32<sup>T34, S97, S130</sup>-CdK5 | 5.6e+06 | 12 |  |
| DARPP-32<sup>T34, S97, S130</sup>-CdK5 -> DARPP-32<sup>T34, T75, S97, S130</sup> + CdK5 |  | 3 |  |
| DARPP-32<sup>S97</sup> + CK1 -> DARPP-32<sup>S97</sup>-CK1 | 4.4e+06 | 12 |  |
| DARPP-32<sup>S97</sup>-CK1 -> DARPP-32<sup>S97, S130</sup> + CK1 |  | 3 |  |
| DARPP-32<sup>T34,S97</sup> + CK1 -> DARPP-32<sup>T34,S97</sup>-CK1 | 4.4e+06 | 12 |  |
| DARPP-32<sup>T34,S97</sup>-CK1 -> DARPP-32<sup>T34,S97,S130</sup> + CK1 |  | 3 |  |
| DARPP-32<sup>T75,S97</sup> + CK1 -> DARPP-32<sup>T75,S97</sup>-CK1 | 4.4e+06 | 12 |  |
| DARPP-32<sup>T75,S97</sup>-CK1 -> DARPP-32<sup>T75,S97,S130</sup> + CK1 |  | 3 |  |
| DARPP-32<sup>T34,T75,S97</sup> + CK1 -> DARPP-32<sup>T34,T75,S97</sup>-CK1 | 4.4e+06 | 12 |  |
| DARPP-32<sup>T34,T75,S97</sup>-CK1 -> DARPP-32<sup>T34,T75,S97,S130</sup> + CK1 |  | 3 |  |
| DARPP-32<sup>T34,S97</sup> + PP2B -> DARPP-32<sup>T34,S97</sup>-PP2B | 7.5e+04 | 120 |  |
| DARPP-32<sup>T34,S97</sup>-PP2B -> DARPP-32<sup>S97</sup> + PP2B |  | 0.03 |  |
| DARPP-32<sup>T34,T75,S97</sup> + PP2B -> DARPP-32<sup>T34,T75,S97</sup>-PP2B | 7.5e+04 | 120 |  |
| DARPP-32<sup>T34,T75,S97</sup>-PP2B -> DARPP-32<sup>T75,S97</sup> + PP2B |  | 0.03 |  |
| DARPP-32<sup>T34,S97,S130</sup> + PP2B -> DARPP-32<sup>T34,S97,S130</sup>-PP2B | 7.5e+04 | 120 |  |
| DARPP-32<sup>T34,S97,S130</sup>-PP2B -> DARPP-32<sup>S97,S130</sup> + PP2B |  | 0.03 |  |
| DARPP-32<sup>T34,T75,S130</sup> + PP2B -> DARPP-32<sup>T34,T75,S130</sup>-PP2B | 7.5e+04 | 120 |  |
| DARPP-32<sup>T34,T75,S130</sup>-PP2B -> DARPP-32<sup>T75,S130</sup> + PP2B |  | 0.03 |  |
| DARPP-32<sup>T34,T75,S97,S130</sup> + PP2B -> DARPP-32<sup>T34,T75,S97,S130</sup>-PP2B | 7.5e+04 | 120 |  |
| DARPP-32<sup>T34,T75,S97,S130</sup>-PP2B -> DARPP-32<sup>T75,S97,S130</sup> + PP2B |  | 0.03 |  |
| DARPP-32<sup>S97,S130</sup> + PP2C -> DARPP-32<sup>S97,S130</sup>-PP2C | 7.5e+06 | 12 |  |
| DARPP-32<sup>S97,S130</sup>-PP2C -> DARPP-32<sup>S97</sup> + PP2C |  | 3 |  |
| DARPP-32<sup>T34,S97,S130</sup> + PP2C -> DARPP-32<sup>T34,S97,S130</sup>-PP2C | 7.5e+06 | 12 |  |
| DARPP-32<sup>T34,S97,S130</sup>-PP2C -> DARPP-32<sup>T34,S97</sup> + PP2C |  | 3 |  |
| DARPP-32<sup>T75,S97,S130</sup> + PP2C -> DARPP-32<sup>T75,S97,S130</sup>-PP2C | 7.5e+06 | 12 |  |
| DARPP-32<sup>T75,S97,S130</sup>-PP2C -> DARPP-32<sup>T75,S97</sup> + PP2C |  | 3 |  |
| DARPP-32<sup>T34,T75,S97,S130</sup> + PP2C -> DARPP-32<sup>T34,T75,S97,S130</sup>-PP2C | 7.5e+06 | 12 |  |
| DARPP-32<sup>T34,T75,S97,S130</sup>-PP2C -> DARPP-32<sup>T34,T75,S97</sup> + PP2C |  | 3 |  |

**Figure 8.**
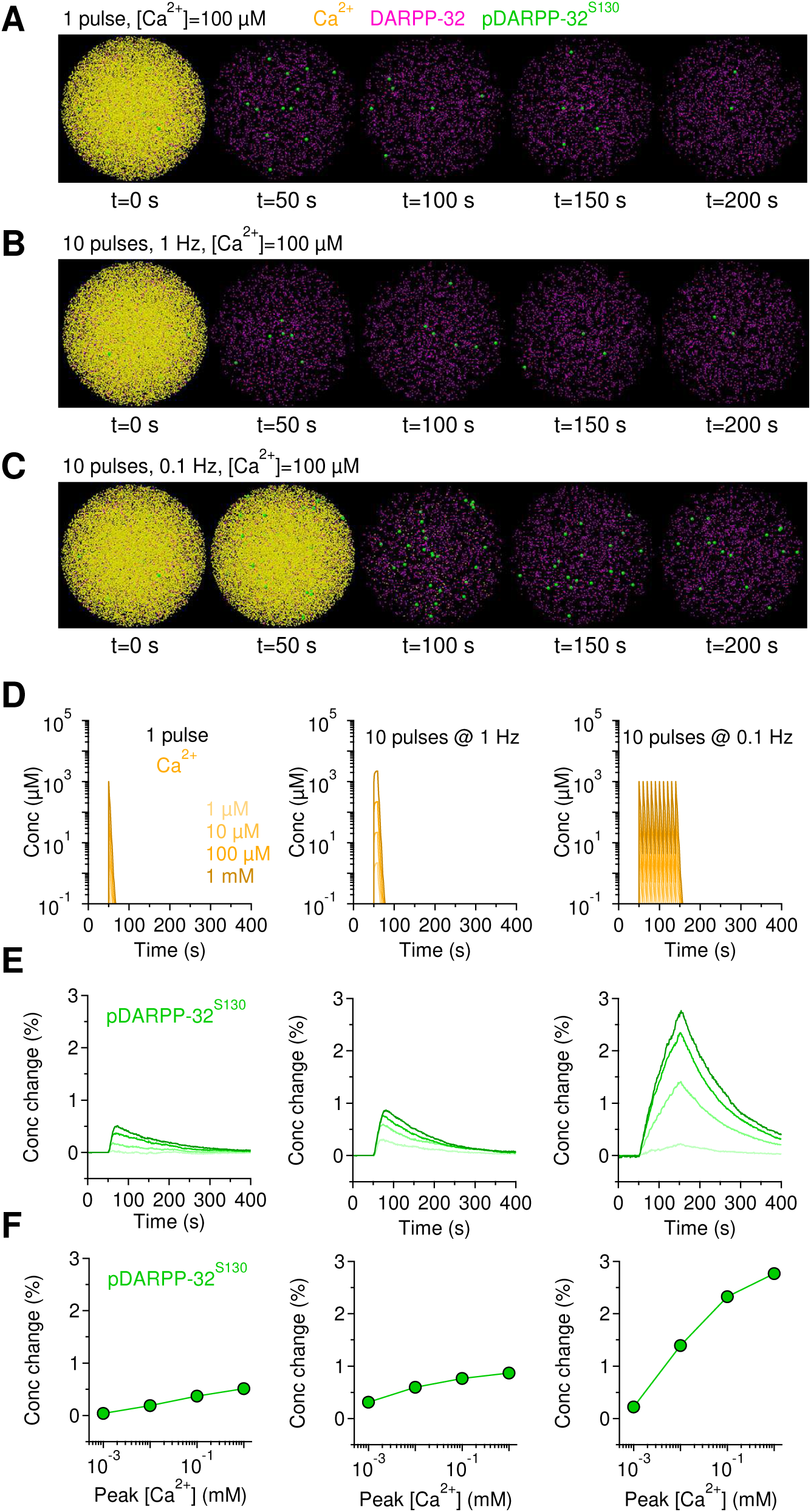
Modeling the effect of Gq signaling cascade activation on DARPP-32 phosphorylation. **(A-C)**Snapshots showing the effect of different patterns of intracellular calcium rise (yellow) on the total concentration of DARPP-32 (magenta) and of pDARPP-32^S130^ (green). **(D)** Time course of intracellular calcium concentration evoked by single (left) or 10 pulse trains of intracellular calcium rise at 1 Hz (middle) and 0.1 Hz (right). Darker colors correspond to higher intracellular calcium concentrations. **(E)** Relative change in intracellular pDARPP-32^S130^. Darker colors correspond to the changes evoked by higher intracellular calcium concentration. **(F)** Summary graph of the change in the peak pDARPP-32^S130^ concentration evoked by increasing concentrations (x-axis) and different patterns of intracellular calcium rise (left to right). Transient increase in the intracellular calcium concentration cause a substantial increase in the intracellular pDARPP-32^S130^ concentration.

## Experimental Design and Statistical Analysis

Data are presented as mean±S.E.M, unless otherwise stated. All experiments were performed on multiple mice of either sex. Statistical significance was determined by Student’s paired or unpaired t-test, as appropriate (IgorPro 6.36). Differences were considered significant at *p*<0.05 (\**p*<0.05; \*\**p*<0.01; \*\*\**p*<0.001).

## RESULTS

### The behavioral phenotype of EAAC1^−/−^ mice

As a first step in our analysis, we performed a comprehensive behavioral analysis of constitutive EAAC1 knock-out mice (EAAC1^−/−^). These mice, originally developed by (Peghini et al., 1997), were obtained by inserting a *pgk* neomycin resistance cassette in the *Slc1a1* gene, which encodes EAAC1 (Peghini et al., 1997). We assessed their general phenotype and tested for the presence of traits that might be indicative of compulsive behaviors. In the SHIRPA screen, we assessed the presence of disturbances in gait, posture, muscle tone deficits and abnormalities in motor control and coordination (**Fig. 1**). We performed this analysis on mice aged P14-35, to cover a wide range of ages starting when EAAC1 reaches its peak expression in the striatum at P14 (Furuta et al., 1997) and ending two weeks after glutamatergic cortico-striatal synapses and dopamine receptors and terminals complete their maturation at P21 (Hattori and McGeer, 1973; Stamford, 1989; Teicher et al., 1995; Sharpe and Tepper, 1998). The results of the SHIRPA screen evidenced subtle motor deficits that had not been reported when the mice were first developed and characterized (Peghini et al., 1997). These subtle abnormalities were present throughout the entire P14-35 age range and could be detected when pooling data from EAAC1^−/−^ mice of either sex (**Fig. 1**) and when analyzing male and female mice separately (**Supplementary Fig. 1-1**).

**Figure 1.**
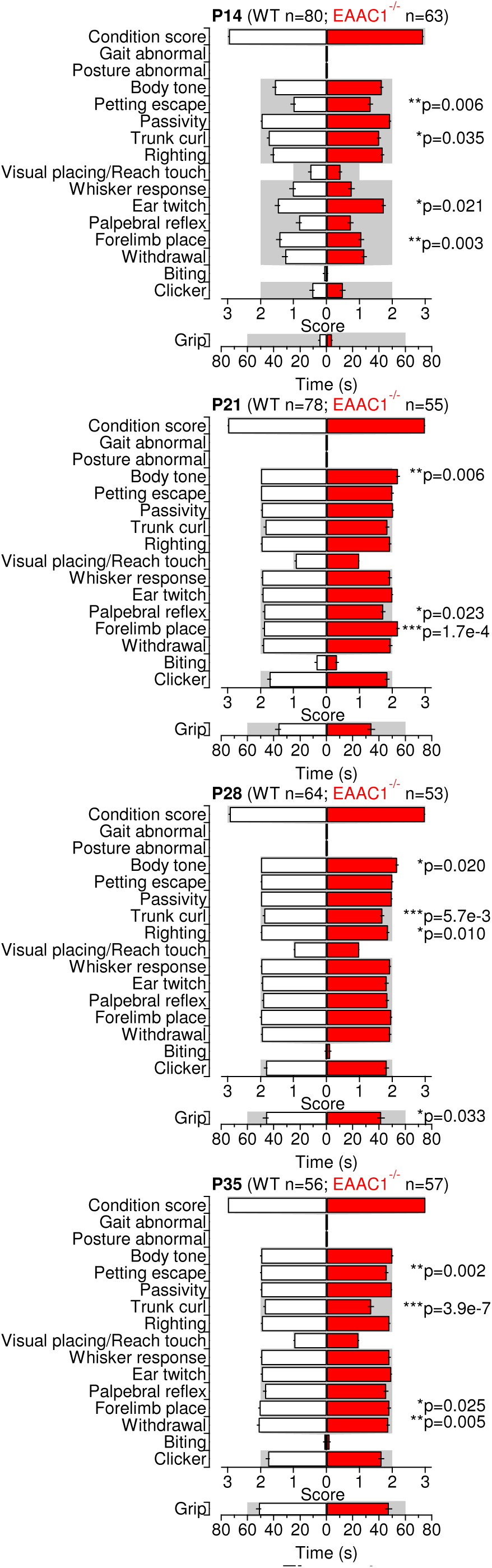
A battery of SHIRPA tests reveals subtle abnormalities in male and female EAAC1^−/−^ mice. **(A)** Parameter scores for a battery of SHIRPA primary screen test in male and female WT (white bars) and EAAC1^−/−^ mice (red bars) aged P14-35. **(B)** As in (A), for male mice.

The overall level of motor activity, measured as distance travelled and time spent on a running wheel, was similar in WT and EAAC1^−/−^ mice (p=0.35 and p=0.12, respectively; **Fig. 2A**). Consistent with these findings, the total distance travelled by WT and EAAC1^−/−^ mice in an open field test was also similar (p=0.28; **Fig. 2B, middle**). What we noticed while performing this test, however, was that the EAAC1^−/−^ mice were less immobile (**p=4.6e-3), because they fidgeted when they resided in a given spot of the open field arena (**Fig. 2B, right**). This restless behavior can be indicative of increased anxiety-like and motor behaviors in EAAC1^−/−^ mice. This hypothesis was supported by data collected using the elevated plus maze test. EAAC1^−/−^ mice showed an increased proportion of entries in the closed arms (**p=0.006; **Fig. 2C**) and spent more time here than in the open arms (*p=0.019; **Fig. 2D**). These results suggest that EAAC1^−/−^ mice show increased anxiety-like behaviors. Consistent with these findings, EAAC1^−/−^ mice buried more marbles than age-matched WT mice in the marble burying test (***p=3.9e-11; **Fig. 2E**), another test used to detect anxious behaviors in mice (Angoa-Perez et al., 2013). The different behavior of WT and EAAC1^−/−^ mice could be detected across the entire age range of mice that we tested (**p-5.5e-3; **Fig. 2F**). Similar results to the ones described in **Fig. 2** were also observed when analyzing male and female mice separately (**Supplementary Fig. 2-1, 2-2**). Together, the results described so far, which were collected using different and complementary behavioral tests, indicate that EAAC1^−/−^ mice are more anxious than WT mice. However, none of these tests points to the presence of specific abnormalities in behaviors that are controlled specifically by the striatum.

**Figure 2.**
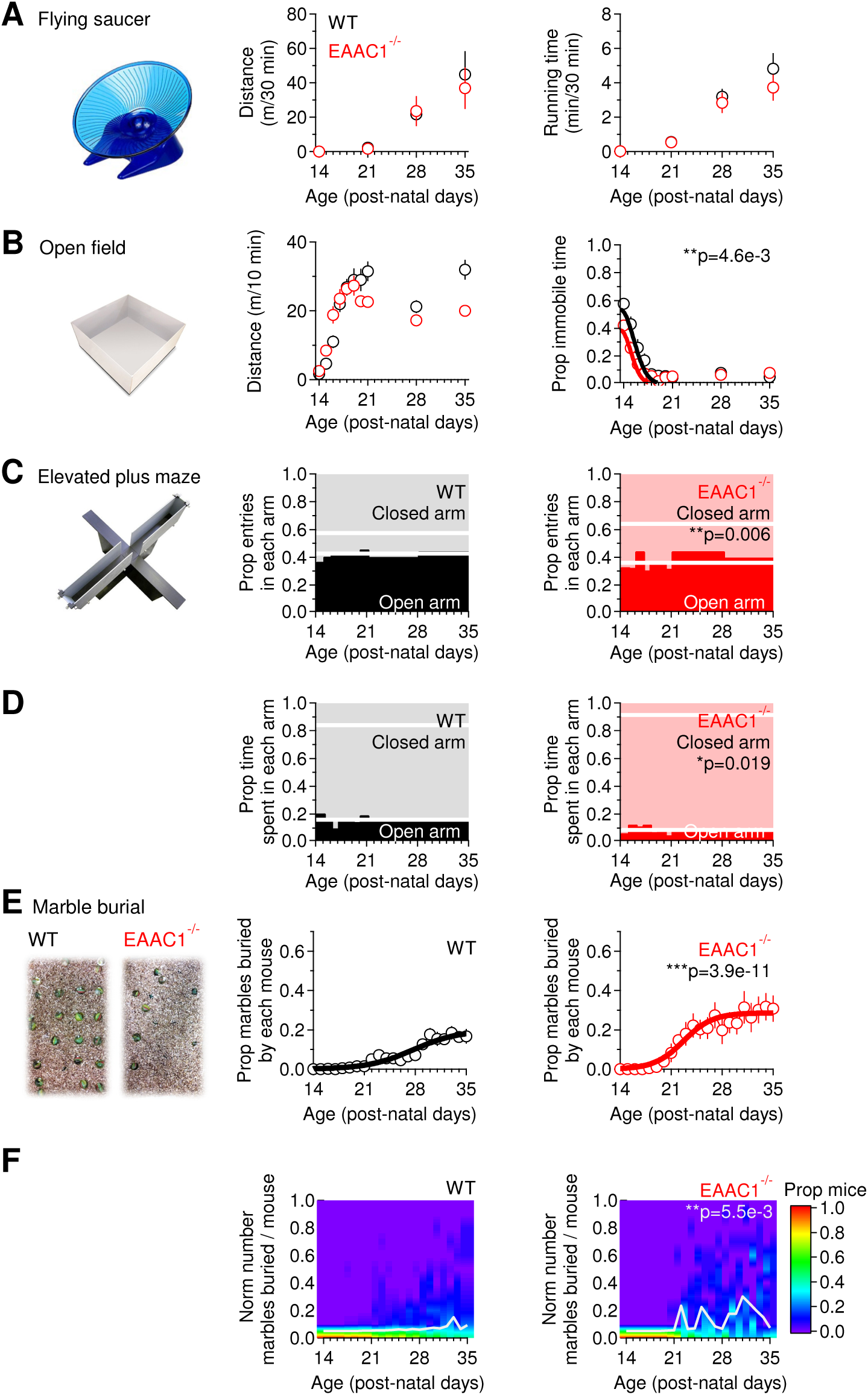
EAAC1’^7^’ mice show similar levels of motor activity but increased anxiety-like behaviors in comparison to WT mice. **(A)** Summary of spontaneous locomotor activity in a free-spinning flying saucer (left), in WT (n=137) and EAAC1^−/−^ mice (n=150) aged P14-35. No significant difference was detected in the mean distance (middle) and running time (right) between WT and EAAC1^−/−^ mice (p=0.35 and p=0.12, respectively). **(B)** In the open field test (left), WT (n=153) and EAAC1^−/−^ mice (n=149) travelled the same distance (p=0.28; middle) but EAAC1^−/−^ mice showed a significant decrease in the proportion of immobile time (**p=4.6e-3; right). **(C)** In the elevated plus maze (left), EAAC1^−/−^ mice (n=90) showed a larger proportion of entries in closed arms in comparison to WT mice (n=118, **p=0.006; left). The thick white lines represent the mean proportion of entries in each arm. **(D)** EAAC1^−/−^ mice spent a larger proportion of time in the closed arms than WT mice (*p=0.019; right). **(E)** In the marble burial test (left), EAAC1^−/−^ mice (n=15; right) dug a significantly larger proportion of marbles than WT mice (n=36, middle; ***p=3.9e-11). **(F)** The color-coding represents the proportion of mice digging a given proportion of marbles (y-axis). The white line describes the behavior of 50% of the mouse population. There is an average increase in the proportion of EAAC1^−/−^ mice digging a larger proportion of marbles (**p=5.5e-3). See **Supplementary Fig. 2-1** and **2-2** for separate analysis of female and male mice.

The striatum exerts a unique role in the control of rule-governed sequential behaviors, including sequential patterned strokes performed during grooming, which are reminiscent of the ritual handwashing behaviors observed in patients with OCD (Berridge and Whishaw, 1992; Kalueff et al., 2016). Interestingly, the striatum is also one of the brain regions that shows hyperactivity in patients with OCD and that has the highest expression levels of EAAC1. EAAC1^−/−^ mice groomed more frequently than WT mice (WT: 6.7±0.7e-3 Hz (n=23), EAAC1^−/−^: 11.3±0.8e-3 Hz (n=42), ***p=4.9e-5), but the duration of each grooming episode was similar to that of WT mice (WT: 55.2±6.7 s (n=42), EAAC1^−/−^: 45.4±3.5 s (n=72), p=0.200; **Fig. 3A-D**). This result could not be attributed to the use of behavioral arenas with transparent floors (see Methods), because the grooming frequency of EAAC1^−/−^ was higher than that of WT mice even when the mice were monitored using a camera positioned above standard mouse cages (WT: 7.6±0.5e-3 Hz (n=39), p=0.34; EAAC1^−/−^: 9.5±0.6e-3 Hz (n=89), p=0.07; WT vs EAAC1^−/−^ *p=0.024). The increased grooming frequency in EAAC1^−/−^ mice did not lead to consistent hair loss or skin lesions associated with other pathological conditions, including trichotillomania (Welch et al., 2007; Feusner et al., 2009). When grooming, the paws of EAAC1^−/−^ mice performed shorter trajectories (WT: 153.3±14.3 mm (n=36), EAAC1^−/−^: 47.9±6.0 mm (n=43), ***p=1.6e-6; **Fig. 3E,F**), suggesting that the execution of fine movements performed during grooming is disrupted in these mice.

**Figure 3.**
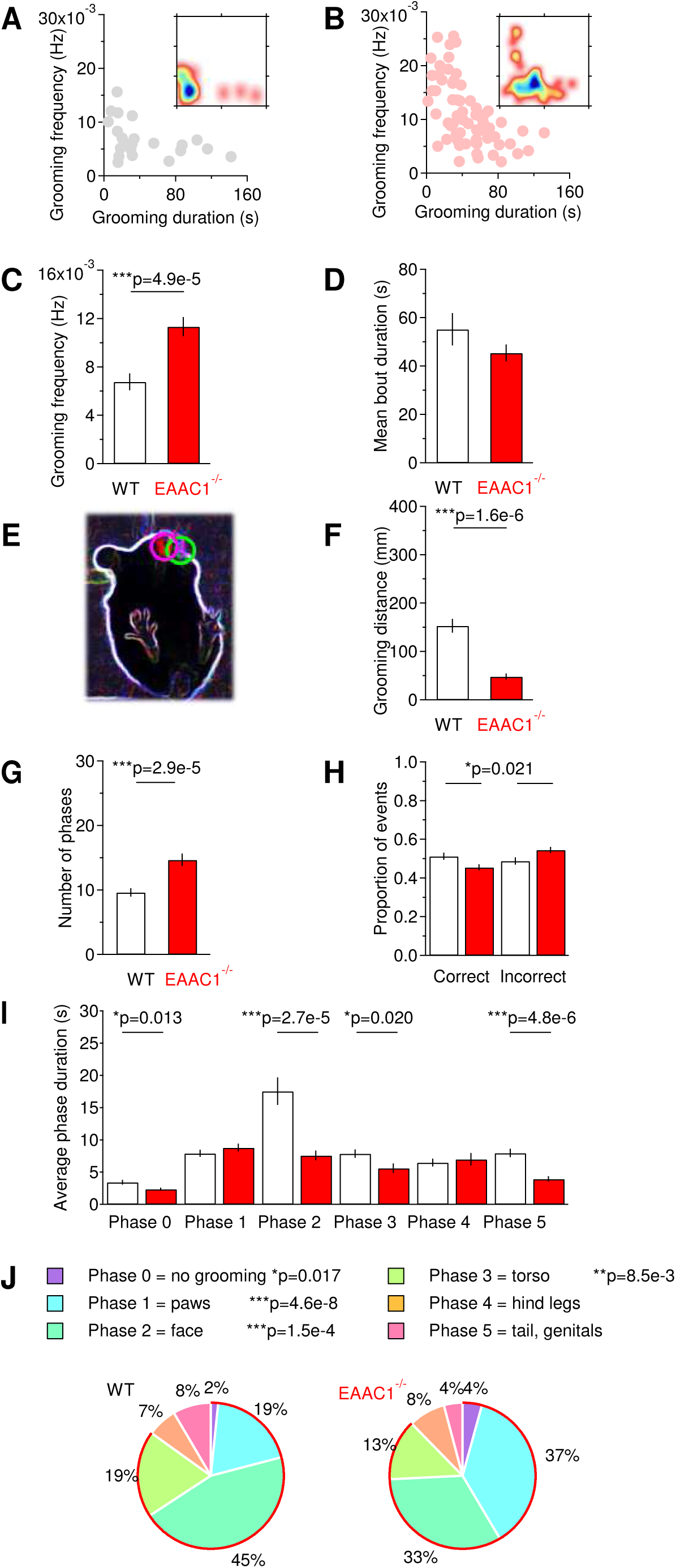
EAAC1’^7^’ mice show disrupted grooming behaviors. **(A)** Relationship between the frequency and duration of grooming episodes in WT mice. The inset represents a density plot of the data, with areas color-coded in blue identify the duration and frequency of the most commonly observed grooming episodes. **(B)** As in (A), for EAAC1^−/−^ mice. **(C)** Summary bar graph of the frequency of grooming episodes (WT (n=23), EAAC1^−/−^ (n=68), ***p=4.9e-5). **(D)** Summary bar graph of the mean duration of individual grooming episodes (WT (n=42), EAAC1^−/−^ (n=72), p=0.200). **(E)** Contour of the mouse body (white) and of the mouse paws (magenta and green circles) detected by the M-Track software, used to perform the analysis of paw trajectories (Reeves et al., 2016). **(F)** Summary bar graph of the mean trajectory length of mice paws as they move to execute a grooming episode (WT (n=36), EAAC1^−/−^ (n=43), ***p=1.6e-6). **(G)** Summary bar graph of the mean number of grooming phases represented in each grooming episode (WT (n=87), EAAC1^−/−^ (n=86), ***p=2.9e-5). **(H)** Proportion of correct (left) and incorrect phase transitions in each grooming episode (WT (n=83), EAAC1^−/−^ (n=85), EAAC1^−/−^p=0.021). **(I)** Summary of the average duration of distinct phases represented in each grooming episode. A decreased representation of phases 0, 2, 3, 5 is detected in EAAC1^−/−^ mice (Phase 0 WT (n=22), EAAC1^−/−^ (n=34), *p=0.013; Phase 1 WT (n=71), EAAC1^−/−^ (n=79), p=0.301; Phase 2 WT (n=73), EAAC1^−/−^ (n=82), ****p=2.7e-5; Phase 3 WT (n=64), EAAC1^−/−^ (n=65), *p=0.020; Phase 4 WT (n=39), EAAC1^−/−^ (n=35), p=0.650; Phase 5 WT (n=30), EAAC1^−/−^ (n=29), ***p=4.8e-6). **(J)** The pie charts provide a representation of the normalized distribution of different phases in each grooming episode. A proportional increase in the representation of phases 0-3 is detected in EAAC1^−/−^ mice. Significant p-values are included in the figure color legend.

Previous work indicates that in mice each grooming episode consists of distinct phases characterized by the execution of specific types of patterned strokes (Kalueff et al., 2007). These different strokes can be classified in six different phases through which mice groom in an orderly sequence along the rostro-caudal axis of their body, from paws to tail (Kalueff et al., 2007). According to this classification, Phase 0 represents time intervals during which mice do not groom. Phases 1-5 describes phases during which mice groom their front paws, face, toes, hind legs and tail/genitals, respectively (Kalueff et al., 2007). The grooming episodes of EAAC1^−/−^ mice contained a larger number of phases than in WT mice (WT: 9.6±0.6 (n=87), EAAC1^−/−^: 14.7±1.0 (n=86), ***p=2.9e-5; **Fig. 3G**). The order of progression from one phase to the next one was disrupted in EAAC1^−/−^ mice, due to the presence of an increased proportion of incorrect transitions (WT: 0.49±0.02 (n=83), EAAC1^−/−^: 0.54±0.01 (n=85), *p=0.021; **Fig. 3H**). EAAC1^−/−^ mice spent less time in Phase 0 (WT: 3.4±0.3 s (n=22), EAAC1^−/−^: 2.3±0.2 (n=34), *p=0.013), Phase 2 (WT: 17.5±2.1 s (n=73), EAAC1^−/−^: 7.6±0.7 s (n=82), ***p=2.7e-5), Phase 3 (WT: 7.9±0.6 s (n=64), EAAC1^−/−^: 5.6±0.7 s (n=65), *p=0.020) and Phase 5 (WT: 8.0±0.7 (n=30), EAAC1^−/−^: 3.9±0.4 s (n=29), ***p=4.8e-6; **Fig. 3I**). WT mice spent most of their grooming time rubbing their torso, whereas EAAC1^−/−^ mice mostly licked their paws (**Fig. 3J**). The increased grooming frequency and the disrupted syntactic structure of grooming episodes point to potential functional abnormalities in the striatum of EAAC1^−/−^ mice and suggest a that EAAC1 might contribute to altered execution of stereotyped motor behaviors in mice.

### EAAC1 shapes synaptic transmission and plasticity in the striatum

We DiRectly tested the hypothesis that EAAC1 controls striatal function *ex vivo* by obtaining extracellular field recordings in acute brain slices containing the DLS, a domain of the striatum specifically implicated with habit learning (Barnes et al., 2005; Yin et al., 2005; Yin et al., 2006). We stimulated excitatory synaptic afferents by positioning a bipolar stimulating electrode in the DLS, 100-200 μm from the recording electrode (**Fig. 4A**) and added the GABA_a_ receptor blocker picrotoxin (100 μM) to the extracellular solution to isolate excitatory from inhibitory responses. The extracellular recordings consist of a short-latency fiber volley (FV) followed by a longer lasting population spike (PS), as described by (Misgeld et al., 1979). As a first step, we confirmed that: *(1)* blocking action potential propagation with the voltage-gated sodium channel blocker TTX (1 μM) abolished both the FV and PS (**Fig. 4B**); *(2)* the GluA antagonist NBQX (10 μM) blocked the PS but not the FV (**Fig. 4C**); *(3)* TTX, applied in the presence of NBQX, blocked the FV (**Fig. 4C**).

In the hippocampus, EAAC1 limits extra-synaptic GluN activation (Scimemi et al., 2009). To test for a similar role of EAAC1 in the DLS, we monitored the effect of the competitive, high-affinity GluN antagonist APV (50 μM) on the field recordings (**Fig. 4D,E**). APV did not induce a significant change in the FV (Norm FV amp WT: 1.03±0.08 (n=12), EAAC1^−/−^: 1.10±0.05 (n=9), p=0.41; **Fig. 4D,E**) and PS/FV amplitude ratio (Norm PS/FV amp WT: 0.99±0.09 (n=12), EAAC1^−/−^: 1.02±0.08 (n=9), p=0.82; **Fig. 4D,E**), consistent with the limited involvement of GluN receptors in mediating synaptic transmission at excitatory synapses in the striatum (Sung et al., 2001b). We performed additional control experiments to determine whether this result could be due to compensatory up-regulation of glial glutamate transporters in the absence of EAAC1, which could also limit GluN activation. However, we did not find any significant difference in the protein expression level of glial glutamate transporters GLAST (Norm GLAST EAAC1^−/−^/WT: 1.02±0.09 (n=10), p=0.81) and GLT-1 (Norm GLT-1 EAAC1^−/−^/WT: 1.07±0.23 (n=8), p=0.77) in protein extracts from the striatum of WT and EAAC1^−/−^ mice (data not shown).

The results of the electrophysiology experiments are important because they suggest that the mechanisms by which EAAC1 controls excitatory transmission in the DLS may be different than in the hippocampus. Other types of glutamate receptors, like the metabotropic group I receptors (mGluRI), are expressed at high levels in extra-synaptic regions along the plasma membrane of striatal neurons (Paquet and Smith, 2003). These receptors are known to modulate cell excitability through a variety of mechanisms including suppression of potassium channels (e.g. I_AHP_, I_M_, I_leak_ and l_slow_) (Charpak et al., 1990; Womble and Moises, 1994; Ikeda et al., 1995; Luthi et al., 1997) and calcium channels (Crepel et al., 1994; Kammermeier et al., 2000). Although blocking mGluRI with type 1 and 5 mGluR antagonists did not affect the amplitude of the FV and PS in WT mice (Norm FV amp WT: 1.08±0.05 (n=13), EAAC1^−/−^: 1.06±0.08 (n=16), p=0.85; **Fig. 4F,G**), it significantly increased the PS/FV amplitude ratio in EAAC1^−/−^ mice (Norm FV amp WT: 0.91±0.09 (n=13), EAAC1^−/−^: 1.21±0.06 (n=16), **p=3.4e-3, WT vs EAAC1^−/−^ *p=0.012; **Fig. 4F,G**). We used separate control experiments to rule out that these results could be due to time-dependent changes in the FV and PS/FV amplitude over the course of our experiments (**Fig. 4H,I**). To do this, we obtained extracellular recordings without adding any drug apart from picrotoxin to the external solution. There was no significant time-dependent change in the FV amplitude (Norm FV amp WT: 0.99±0.06 (n=8), p=0.93; Norm FV amp EAAC1^−/−^: 1.12±0.09 (n=7), p=0.24; WT vs EAAC1^−/−^ p=0.28) and in the PS/FV amplitude ratio, in WT and EAAC1^−/−^ mice (Norm PS/FV amp WT: 1.16±0.07 (n=8), p=0.058; Norm PS/FV amp EAAC1^−/−^: 1.16±0.17 (n=7), p=0.38; WT vs EAAC1^−/−^ p=1.00; **Fig. 4H,I**). This suggests that the different sensitivity to mGluRI antagonists of the PS/FV ratio in EAAC1^−/−^ mice is genuine. By obtaining input-output curves, we confirmed that this effect could be detected over a broad range of stimulus intensities in EAAC1^−/−^(n=16), not in WT mice (n=13, **p=2.1e-3; **Fig. 4J**).

The increased PS/FV sensitivity to mGluRI antagonists could be explained by increased mGluRI expression or by increased extracellular glutamate concentration in EAAC1^−/−^ mice. Western blot analysis showed that the mGluRI protein expression level is similar in WT and EAAC1^−/−^ mice (Norm band intensity WT: 1±0.13 (n=6), EAAC1^−/−^: 1.23±0.23 (n=8), p=0.41; **Fig. 4K**). To determine whether there could be an increase in the extracellular glutamate concentration in the absence of EAAC1, we measured tonic GluN currents. Briefly, we voltage clamped MSNs at -70 mV in the presence of Mg^2+^-free external solution containing blockers of GABA_A_ (picrotoxin, 100 μM) and GluA receptors (NBQX, 10 μM) and measured the change in the holding current evoked by blocking GluN receptors with APV (50 μM). We did not detect any significant change in the tonic GluN current (WT: 10.1±4.1 pA (n=6), EAAC1^−/−^: 15.8±9.3 pA (n=6), p=0.59) and tonic GluN current density (tonic current/cell capacitance) in WT and EAAC1^−/−^ mice (WT: 0.55±0.43 pA/pF (n=6), EAAC1^−/−^: 0.36±0.19 pA/pF (n=6), p=0.70). Is this result consistent with our knowledge of mGluRI kinetics of activation and of glutamate transporter control of ambient glutamate concentration? We addressed this question using a modeling approach. *First*, we estimated the mGluRI open probability over a broad range of extracellular glutamate concentrations using a kinetic model of mGluRI ((Marcaggi et al., 2009); **Fig. 4L**). *Second*, we calculated the mGluRI open probability at the experimentally-measured extracellular glutamate concentration in the striatum (∼25 nM; (Chiu and Jahr, 2017)) and the estimated concentration of glutamate transporters (∼140 μM; (Lehre and Danbolt, 1998)). *Third*, we used steady-state equations to determine the effect of a progressive reduction in the glutamate transporter concentration on the extracellular glutamate concentration and the mGluRI open probability. The results show that the mGluRI open probability is very low (P_o_∼6e-4) when the glutamate transporter concentration is 140 μM (i.s. Norm [Transporter=1]). Reducing the glutamate transporter concentration by 5%, consistent with the expected change in glutamate transporter concentration in the absence of EAAC1 (Danbolt, 2001), causes ~10 nM increase in the ambient glutamate concentration (**Fig. 4L**). This increase in ambient glutamate concentration does not cause a significant change in the mGluRI open probability. Therefore, the results of the Western blot analysis and the modeling suggest that the increased contribution of mGluRI to the PS/FV ratio in EAAC1^−/−^ mice is not due to either increased mGluRI expression or increased tonic mGluRI activation. Instead, it is consistent with increased phasic activation of mGluRI in the absence of EAAC1.

The effect of EAAC1 on mGluRI activation is noteworthy because the activation of different types of glutamate receptors in the DLS is crucial for the expression of long-term plasticity. Here, long-term potentiation (LTP) relies on GluN and D1R activation (Calabresi et al., 1992a; Partridge et al., 2000; Spencer and Murphy, 2000). Long-term depression (LTD) relies on mGluRI and D2R activation and on post-synaptic calcium influx via L-type calcium channels (Calabresi et al., 1992b; Gubellini et al., 2001; Sung et al., 2001a). In our experiments, high-frequency stimulation (HFS; 100 Hz × 1 s) of the DLS evoked LTD or LTP, depending on the extracellular calcium concentration ([Ca^2+^]_o_; **Fig. 5A**). In WT mice, HFS induced LTP at a calcium concentration that mimics the one found in the cerebrospinal fluid ([Ca^2+^]_o_=1.2 mM; Norm PS/FV amp WT: 1.21±0.09 (n=7), *p=0.047). HFS did not induce long-term plasticity at [Ca^2+^]_o_=2.5 mM (Norm PS/FV amp WT: 1.09±0.07 (n=7), p=0.22). Increasing the extracellular calcium concentration to *[Ca^2+^]_o_=* 3.5 mM led to a robust LTD, likely because of the increased driving force for post-synaptic calcium influx (Norm PS/FV amp WT: 0.84±0.06 (n=11), *p=0.032; **Fig. 5A,C**). In EAAC1^−/−^ mice, HFS did not induce any form of plasticity at any of the tested extracellular calcium concentrations (Norm PS/FV amp [Ca^2+^]_o_=1.2 mM EAAC1^−/−^: 1.04±0.06 (n=8), p=0.50; [Ca^2+^]_o_=2.5 mM EAAC1^−/−^: 0.96±0.04 (n=12), p=0.28; [Ca^2+^]_o_=3.5 mM EAAC1^−/−^: 1.05±0.06 (n=7), p=0.46; **Fig. 5B,D**). Loss of LTD in EAAC1^−/−^ mice is consistent with altered mGluRI activation. However, loss of LTP suggests that other receptors, in addition to mGluRI, might be disrupted in the absence of EAAC1. Therefore, we performed additional experiments to determine what molecular mechanism could account for the loss of LTP in the DLS of EAAC1^−/−^ mice.

**Figure 5.**
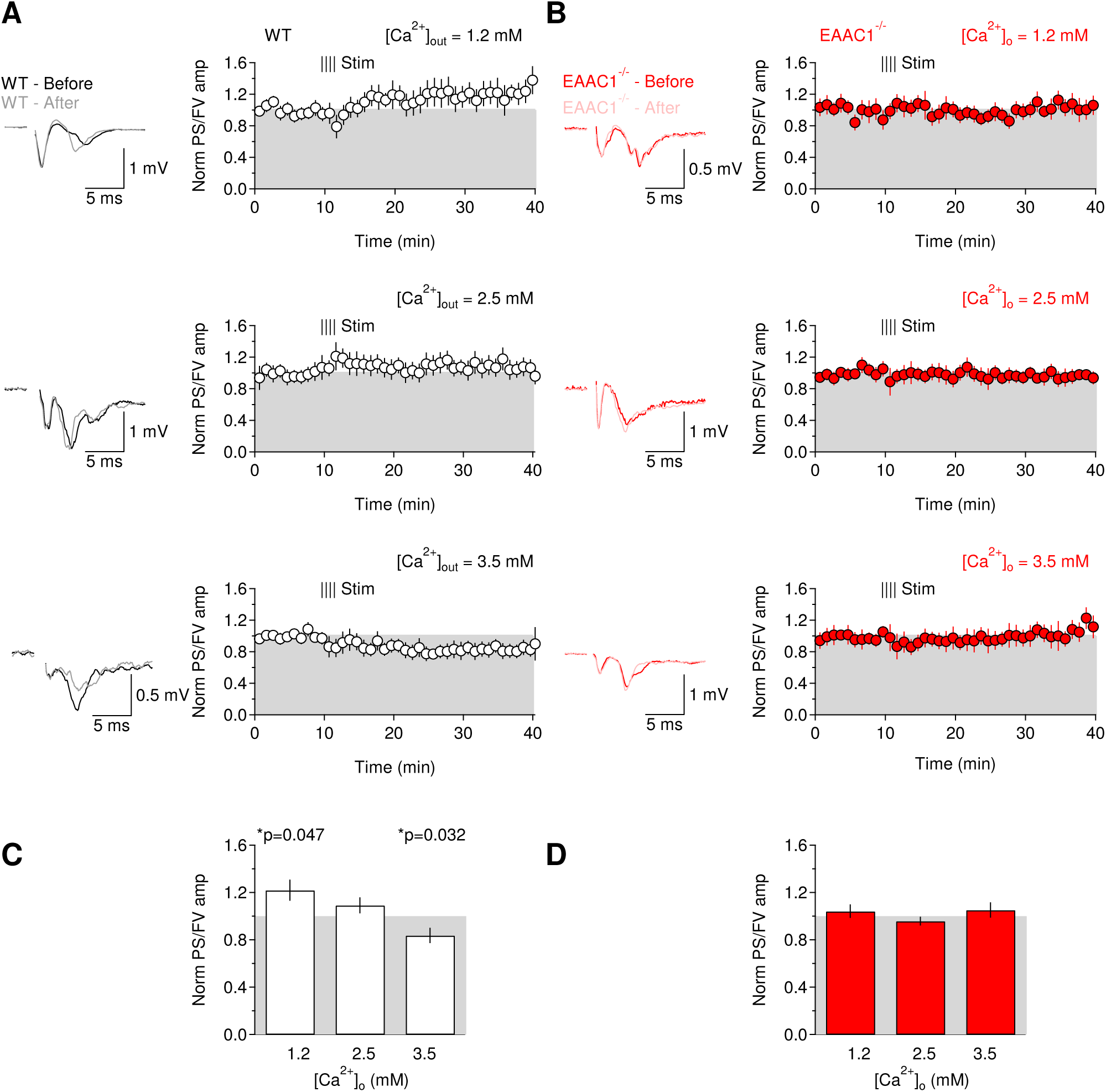
The Ca^2+^-dependence of long-term plasticity is lost in EAAC1^−/−^ mice. **(A)** Left: extracellular recordings obtained 5 min before (black trace) and 30 min after applying a high-frequency stimulation protocol (HFS: 100 Hz, 1 s) to the DLS of WT mice (grey trace). The recordings were obtained in the presence of extracellular solutions containing [Ca^2+^]=1.2 mM (top panel), [Ca^2+^]=2.5 mM (middle panel) and [Ca^2+^]=3.5 mM (bottom panel). Each trace represents the average of 20 consecutive sweeps. The shaded area represents the S.E.M. Right: time course of baseline-normalized field recordings. Each symbol represents the average of three consecutive time points. The notation “Norm PS/FV” on the y-axis refers to the amplitude ratio of the population spike and fiber volley normalized by the one measured before applying the high-frequency stimulation protocol. **(B)** As in (A), for EAAC1^−/−^ mice. **(C)** Summarized effect of HFS on the PS/FV ratio in WT mice at three different extracellular [Ca^2+^] (1.2 mM: (n=7) *p=0.047; 2.5 mM: (n=7) p=0.22; 3.5 mM: (n=11) *p=0.032). **(D)** Summarized effect of HFS on the PS/FV ratio in EAAC1^−/−^ mice at three different extracellular [Ca^2+^] (1.2 mM: (n=8) p=0.50; 2.5 mM: (n=12) p=0.28; 3.5 mM: (n=7) p=0.46).

### EAAC1 promotes dopamine receptor expression

Loss of LTP in EAAC1^−/−^ mice is surprising because this form of plasticity in the striatum does not require mGluRI but GluN and D1R activation. The electrophysiology experiments in **Fig. 4** do not support a different contribution of GluN receptors to excitatory synaptic transmission in the DLS of EAAC1^−/−^ mice, and therefore point to an effect of EAAC1 on D1Rs. To test this hypothesis, we measured the mRNA (**Fig. 6**) and protein levels of D1R and D2R (**Fig. 7**) in the striatum of WT and EAAC1^−/−^ mice, using Real-Time quantitative Reverse Transcription PCR (qRT-PCR) and Western blot analysis, respectively. The sensitivity of the qRT-PCR technique was validated by measuring mRNA levels for the *Slc1a1* gene, encoding EAAC1, in the striatum of WT and EAAC1^−/−^ mice (**Fig. 6A-D**). We confirmed that the mRNA level of EAAC1 was negligible in EAAC1^−/−^ mice (**Fig. 6A-C;** Norm *Slc1a1* EAAC1^−/−^/WT: 0.05±0.01 (n=5), ***p=1.5e-7) and that the mRNA levels of EAAC1 were similar in the DLS and VMS (p=0.42; **Fig. 6D**). The mRNA level of D1R in the entire striatum, not D2R, was significantly reduced in EAAC1^−/−^ compared to WT mice (Norm *Drd1a* EAAC1^−/−^/WT: 0.65±0.10 (n=11), **p=5.0e-3; Norm *Drd2* EAAC1'^7^'/WT: 0.89±0.14 (n=11), p=0.48; **Fig. 6E-G**). Similar results were obtained when we analyzed the DLS and VMS separately, for D1R (Norm DLS *Drd1a* EAAC1^−/−^/WT: 0.82±0.06 (n=6), *p=0.037; Norm VMS *Drd1a* EAAC1^−/−^/WT: 0.53±0.08 (n=6), **p=1.6e-3; **Fig. 6H-J**) and D2R (Norm DLS *Drd2* EAAC1^−/−^/WT: 1.15±0.33 (n=6), p=0.32; Norm VMS *Drd2* EAAC1^−/−^/WT: 0.79±0.29 (n=6), p=0.14; **Fig. 6K-M**), suggesting that they are widespread throughout the entire striatal formation.

Changes in D1R mRNA might be associated with altered D1R protein expression, which we tested using Western blot analysis. As a first step, we validated the sensitivity and specificity of the antibodies directed against D1R and D2R. *First*, we performed immuno-labeling experiments on striatal sections from mice in which the genetically encoded red fluorescent protein (RFP) mCherry was selectively expressed either in D1- or D2-MSNs (**Supplementary Fig. 7-1**). In the DLS, the anti-D1R antibody labelled a significant proportion of RFP-expressing cells in sections from D1^Cre/+^:Ai9^T8/0^ mice (0.70±0.08 (n=4)) and a very small proportion of RFP-expressing cells in sections from A2A^Cre/+^:Ai9^Tg/0^ mice (0.09±0.03 (n=3); **Supplementary Fig. 7-1A**). The vast majority of immuno-labelled cells were RFP-expressing D1-MSNs (0.84±0.04 (n=4)), not D2-MSNs (0.13±0.03 (n=3)), **Supplementary Fig. 7-1A**). Likewise, the anti-D2R antibody labelled a significant proportion of RFP-expressing cells in sections from A2A^Cre/+^:Ai9^Tg/0^ mice (0.83±0.07 (n=4)) and a very small proportion of RFP-expressing cells in sections from D1^Cre/+^:Ai9^Tg/0^ mice (0.08±0.04 (n=3); **Supplementary Fig. 7-1B**). The vast majority of immuno-labelled cells were RFP-expressing D2-MSNS (0.89±0.07 (n=4)), not D1-MSNs (0.14±0.05 (n=3)), **Supplementary Fig.7-1B**). These data showed that the anti-D1R/D2R antibodies are sensitive and specific. To further validate their specificity in Western blot analysis, we compared the D1R protein expression in the striatum and in the cortex, where D1R and D2R are less abundantly expressed (Gong et al., 2003; Gong et al., 2007). Accordingly, the protein expression of D1R was significantly lower in the cortex compared to the striatum (Norm D1R striatum: 1.00±0.14 (n=8), Norm D1R cortex: 0.60±0.08 (n=8), *p=0.037; **Supplementary Fig. 7-1C**). Similar results were obtained when using the anti-D2R antibody (Norm D2R striatum: 1.00±0.10 (n=14), Norm D2R cortex: 0.69±0.08 (n=16), *p=0.022; **Supplementary Fig. 7-1D**). If the antibodies used for these experiments specifically labelled D1R or D2R bands, these bands should no longer be detected after pre-incubating the antibodies with their corresponding control peptide antigen. Consistent with this hypothesis, no band was detected by either antibody in pre-adsorption experiments (Norm D1R peptide: 0.07±0.01 (n=7), ***p=3.7e-4, Norm D2R peptide: 0.12±0.07 (n=7), ***p=1.1e-6; **Supplementary Fig. 7-1C,D**). Having confirmed the specificity of our antibodies, we asked whether the reduced D1R mRNA levels were associated with altered D1R protein levels in EAAC1^−/−^ mice. Consistent with the qRT-PCR data, we detected lower levels of D1R in protein extracts from the entire striatum of EAAC1^−/−^ mice, while no change in D2R expression was detected (Norm D1R EAAC1'VwT: 0.69±0.03 (n=8), ***p=2.7e-4; Norm D2R EAAC1^−/−^/WT: 0.86±0.16 (n=8), p=0.51; **Fig. 7A**).

To determine whether the reduced of D1R protein expression could be consequent to loss of D1-MSNs in EAAC1^−/−^ mice, we compared the proportion and density of RFP-expressing cells in D1^Cre/+^:Ai9^Tg/0^ and D1^Cre/+^:Ai9^Tg/0^EAAC1^−/−^ mice, D1^tdTomato/+^ and D1^tdTomato/+^: EAAC1^−/−^ mice, A2A^Cre/+^:Ai9^Tg/0^ and A2A^Cre/+^:Ai9^Tg/0^:EAAC1^−/−^ mice (**Supplementary Fig. 7-2**). The proportion of RFP-expressing cells was calculated as the ratio between red cells (expressing RFPs) and blue cells (labelled with DAPI), whereas the density of RFP-expressing cells was calculated as the number of red cells in the matrix region of the DLS. There was no significant difference in the density of RFP-expressing D1-MSNs in WT and EAAC1^−/−^ mice, in the DLS (WT: 0.81e-3±7.10e-5 μm^-2^ (n=11), EAAC1^−/−^ 0.83e-3±0.12e-3 μm^-2^ (n=6), p=0.90) and VMS (WT: 0.69e-3±0.11e-3 μm^-2^ (n=6), EAAC1^−/−^ 0.99e-3±8.15e-5 μm^-2^ (n=6), p=0.073). The proportion of D1-MSNs was also similar in WT and EAAC1^−/−^ mice in the DLS (WT: 0.39±0.03 (n=11), EAAC1^−/−^: 0.41±0.03 (n=6), p=0.64) and VMS (WT: 0.31±0.06 (n=7), EAAC1^−/−^: 0.45±0.02 (n=6), p=0.055). Similar results were obtained when measuring the density of RFP-expressing D2-MSNs in the presence and absence of EAAC1, in the DLS (WT: 1.12e-3±3.78e-5 μm^-2^ (n=6), EAAC1^−/−^ 1.07e-3±5.94e-5 μm^-2^ (n=5), p=0.14) and VMS (WT: 0.89e-3±3.38e-5 μm^-2^ (n=6), EAAC1^−/−^ 0.86e-3±2.61e-5 μm^-2^ (n=5), p=0.58). The density of D2-MSNs was larger in the DLS than in the VMS, but this difference was present in both WT (WT DLS: 1.2e-3±3.8e-5 (n=6), VMS: 3.9e-4±3.4e-5 (n=6), **p=1.7e-4) and EAAC1^−/−^ mice (EAAC1^−/−^ DLS: 1.1e-3±5.9e-5 (n=5), VMS: 0.9e-3±2.6e-5 (n=5), *p=0.022) and the proportion of D2-MSNs was also similar in WT and EAAC1^−/−^ mice, in the DLS (WT: 0.40±0.01 (n=6), EAAC1^−/−^: 0.40±0.02 (n=5), p=0.93) and VMS (WT: 0.38±0.02 (n=6), EAAC1^−/−^: 0.40±0.01 (n=5), p=0.31). Together, these results suggest that loss of EAAC1 is not associated with altered density of D1- or D2-MSNs. Therefore the reduced D1R expression in EAAC1^−/−^ mice might be due to altered trafficking and internalization of D1R from the cell membrane.

The expression of dopamine receptors is subject to intracellular regulation by a variety of signaling molecules including the phosphoprotein DARPP-32, which has been suggested to act as a robust integrator of glutamatergic and dopaminergic transmission (Gould and Manji, 2005; Fernandez et al., 2006)). Although the total level of DARPP-32 was similar in WT and EAAC1^−/−^ mice (Norm DARPP-32 EAAC1^−/−^/WT: 1.00±0.05 (n=12), p=0.92; **Fig. 7B**), the ability of this protein to modulate dopaminergic and glutamatergic signaling depends not only on its abundance but on its phosphorylation state (Greengard et al., 1999). The signaling cascades that modulate the phosphorylation state of DARPP-32 include the Gq signaling cascades activated by mGluRI (Liu et al., 2001; Liu et al., 2002; Svenningsson et al., 2004; Nishi et al., 2005). Out of the four known phosphorylation sites of DARPP-32 (T34, T75, S97 and S130), the only one differentially phosphorylated in EAAC1^−/−^ mice was pDARPP-32^S130^ (pDARPP- 32^S130^ WT: 0.28±0.08 (n=14), EAAC1^−/−^: 0.60±0.10 (n=17), *p=0.024; **Fig. 7C**). The almost 3-fold increase in pDARPP-32^S130^ phosphorylation indicates that the phosphorylation pattern of pDARPP-32 is profoundly disrupted in the absence of EAAC1 (Norm pDARPP-32^5130^ EAAC1^−/−^/WT: 2.70±0.44 (n=14), **p=2.1e-3; **Fig. 7D,E**).

Is it plausible that changes in signaling pathways coupled to mGluRI, known to activate phospholipase C (PLC), stimulate the generation of inositol 1,4,5-trisphosphate (IP_3_) and mobilize calcium from internal stores (Casabona et al., 1997), would increase pDARPP-32^S130^ phosphorylation? We addressed this question using a modeling approach based on 3D reaction-diffusion Monte Carlo simulations of calcium diffusion and DARPP-32 phosphorylation (**Fig. 8**). We modelled the geometry of the post-synaptic terminal as a 1 μm^3^ sphere, consistent with the average size of spine heads of striatal neurons (Forlano and Woolley, 2010). At the beginning of each simulation, the sphere contained 3,000 DARPP-32 molecules and a number of protein kinases and phosphatases known to modulate the phosphorylation state of DARPP-32 (**Table 6**). We then allowed the system to equilibrate for 150 s. Under these baseline conditions the resting concentration of pDARPP-32S^130^ was 10 nM. We then released variable amounts of calcium (1 μM-1 mM) from the center of the sphere to simulate the intracellular calcium rise coupled to mGluRI activation. Single pulses or trains of 10 pulses (0.1-10 Hz) evoked a detectable increase pDARPP-32^S130^, suggesting that the altered activation of Gq signaling pathways coupled to mGluRI and the consequent increase in intracellular calcium concentration can alter pDARPP-32^S130^ phosphorylation.

### EAAC1 controls plasticity and dopamine receptor expression via signaling pathways coupled to mGluRI activation

If the loss of synaptic plasticity (**Fig. 5**), reduced D1R expression (**Fig. 6, 7A**), altered phosphorylation of DÂRPP-32 (**Fig. 7B-E**) and disrupted grooming behaviors in EAAC1^−/−^ mice (**Fig. 3**) were all triggered by increased mGluRI activation (**Fig. 4**), we would expect to convert the functional phenotype of EAAC1^−/−^ into that of WT mice by blocking mGluRI. Consistent with this hypothesis, blocking mGluRI with LY367385 (50 μM) and MPEP (10 μM) rescued LTP and LTD in EAAC1^−/−^ mice (Norm PS/FV amp [Ca^2+^]_o_=1.2 mM: 1.28±0.11 (**n=22**), *p=0.015; Norm PS/FV amp [Ca^2+^]_o_=2.5 mM: 1.08±0.08 (**n=12**), p=0.36; Norm PS/FV amp [Ca^2+^]_o_=3.5 mM: 0.84±0.05 (n=8), *p=0.018; **Fig. 9**). This pharmacological treatment also rescued D1R expression (Norm D1R EAAC1^−/−^: 1.13±0.13 (n=9), p=0.35; Norm D2R EAAC1' ^7^': 1.33±0.14 (n=7), p=0.06; **Fig. 10A**) and reduced pDARPP-32^S130^ phosphorylation (Norm pDARPP-32^S130^ EAAC1^−/−^/WT: 0.37±0.11 (n=9), ***p=2.1e-4; **Fig. 10C-E**). These findings are consistent with an effect of mGluRI on pDARPP-32^S130^ phosphorylation (Liu et al., 2001; Liu et al., 2002). However, this broad pharmacological approach also induced a significant reduction in the total level of DARPP-32 (Norm DARPP-32: 0.87±0.04 (n=9), *p=0.029; **Fig. 10B**) and a decrease in pDARPP-32^T75^ phosphorylation (Norm pDARPP-32^T75^ EAAC1^−/−^/WT: 0.45±0.05 (n=12), ***p=1.0e-5; **Fig. 10C-E**). These effects were not detected in EAAC1^−/−^ mice (**Fig. 7**) and may be consequent to the blockade of mGluRI in multiple regions of the brain present in the slice preparations. For this reason, we felt compelled to test more specific approaches to recapitulate more faithfully the molecular and behavioral phenotype of EAAC1^−/−^ mice.

**Figure 9.**
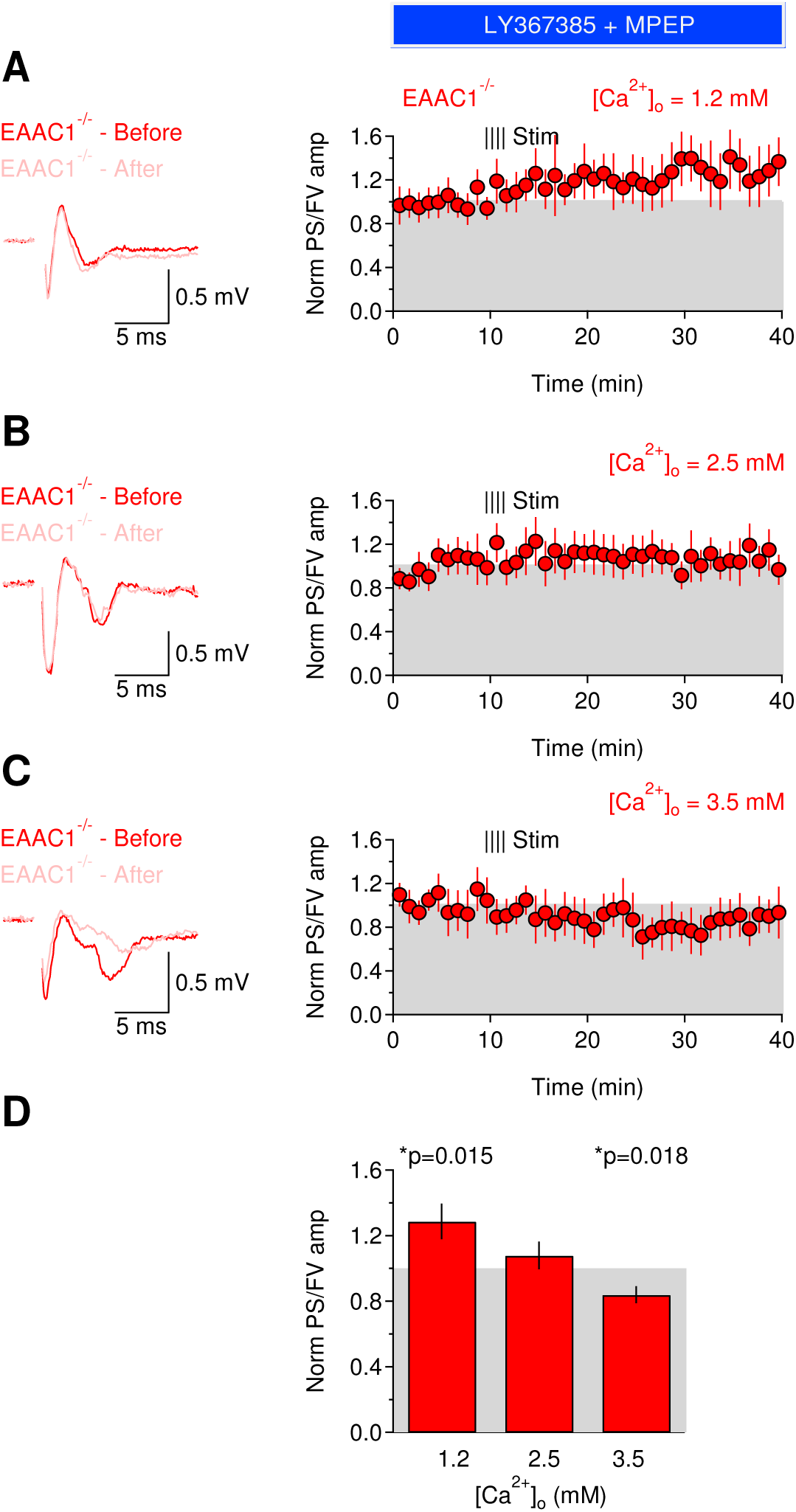
Blocking mGluRI activation rescues the Ca^2+^-dependence of long-term plasticity in EAAC1^−/−^ mice. **(A)** Left: extracellular recordings performed in the presence of the mGluRI blockers LY367385 (50 μM) and MPEP (10 μM) and obtained 5 min before (red trace) and 30 min after applying a high-frequency stimulation protocol (HFS: 100 Hz, 1 s) to the DLS of EAAC1^−/−^ mice (pink trace). The recordings were obtained in the presence of extracellular solutions containing [Ca^2+^]=1.2 mM. Each trace represents the average of 20 consecutive sweeps. The shaded area represents the S.E.M. Right: time course of baseline-normalized field recordings. Each symbol represents the average of three consecutive time points. The notation “Norm PS/FV” on the y-axis refers to the amplitude ratio of the population spike and fiber volley normalized by the one measured before applying the high-frequency stimulation protocol. **(B)** As in (A), in the presence of extracellular solutions containing [Ca^2+^]=2.5 mM. **(C)** As in (A), in the presence of extracellular solutions containing [Ca^2+^]=3.5 mM. **(D)** Summarized effect of HFS on the PS/FV ratio in EAAC1^−/−^ mice in the presence of mGluRI blockers and at three different extracellular [Ca^2+^] (1.2 mM (n=22), *p=0.015; 2.5 mM (n=12), p=0.36; 3.5 mM (n=8), *p=0.018).

### Activation of signaling pathways coupled to mGluRI recapitulates the molecular and behavioral phenotype of EAAC1* mice in WT mice

In the striatum, mGluRI are expressed in both D1- and D2-MSNs (Tallaksen-Greene et al., 1998). The observed reduction of D1R in EAAC1^−/−^ mice, however, suggests a higher sensitivity to mGluRI activation in D1- compared to D2-MSNs. If this were true, we would expect cell-specific activation of G-protein coupled signaling cascades activated by mGluRI in D1-MSNs to mimic more closely the molecular and behavioral phenotype of EAAC1^−/−^ mice. This hypothesis can be tested by using a chemogenetic approach, based on the combined use of BAC transgenic mouse lines and Designer Receptors Exclusively Activated by Designer Drugs (DREADDs) (Armbruster et al., 2007). The conditional expression of DREADDS in D1^Cre/+^ mice allowed us to activate Gq signaling cascades like those coupled to mGluRI exclusively in D1-MSNs. To accomplish this goal, we performed unilateral stereotaxic injections of AAV-hSyn-DIO-hM3D(Gq)-mCherry in the DLS of D1^Cre/+^ mice aged P14-16 (**Fig. 11, left**). We confirmed the successful DREADDs transfection of the injected striatum by measuring the expression of the mCherry protein in the striatum and cortex of the injected and non-injected brain. The non-injected striatum and the adjacent cortex were not expected to express mCherry, and were used as internal controls to confirm the accuracy of the stereotaxic injections in the DLS, not in the adjacent cortex. One week after the surgery, the mice were habituated to daily I/P saline injections, to prevent any bias of our results by acute stress responses to the I/P injections of the DREADD activator clozapine-N-oxide (CNO) two weeks after surgery. DREADD activation is maximal 1-1.5 hours after the CNO injection and starts declining after 3 hours (Gomez et al., 2017; Raper et al., 2017). Therefore, any effect of CNO is likely to happen within this time window. One hour after the CNO injections, we video monitored the mice to measure their grooming activity and then sacrificed them two hours after the CNO injection, to extract the striatum for Western blot analysis (**Fig. 11A**). We then measured the protein levels of mCherry, D1R, D2R and pDARPP-32 and compared their expression levels in the injected and non-injected striatum and cortex (**Fig. 11B-D left**). We detected a substantial increase in mCherry expression only in the injected striatum of D1^Cre/+^ mice, not in the adjacent cortex, confirming the accuracy of our stereotaxic injections (hM3D(Gq) Cortex: 1.22±0.20 (n=10), p=0.28; hM3D(Gq) Striatum: 2.89±0.80 (n=10), *p=0.040; **Fig. 11B, left**). DREADD-mediated activation of Gq signaling cascades led to reduced D1R expression (hM3D(Gq): 0.79±0.07 (n=9), *p=0.013), without affecting D2R expression (hM3D(Gq): 0.91±0.10 (n=8), p=0.35; **Fig. llC, left**). Gq signaling activation also caused a significant and specific increase in the phosphorylation of pDARPP-32^S130^ in D1^Cre/+^ mice (hM3D(Gq): 2.48±0.49 (n=ll), **p=5.6e-3; **Fig. 11D, left**). These results show that activation of Gq signaling cascades like the ones activated by mGluRI can lead to increased phosphorylation of pDARPP-32^S130^ and reduced D1R expression like in EAAC1^−/−^ mice, possibly because of increased D1R internalization.

As an additional test of our hypothesis, we performed similar experiments using stereotaxic injections of AAV-hSyn-DIO-hM3D(Gq)-mCherry in the DLS of A2A^Cre/+^ mice. We confirmed the increase in mCherry expression in the injected striatum and not in the adjacent cortex in A2A^Cre/+^ mice (hM3D(Gq) Cortex: 1.18±0.31 (n=8), p=0.59; hM3D(Gq) Striatum: 2.06±0.52 (n=23), *p=0.035; **Fig. 11B, right**). In this case, DREADD-mediated activation of Gq signaling cascades in D2-MSNs did not alter D1R expression (hM3D(Gq): 1.10±0.15 (n=9), p=0.63) and D2R expression (hM3D(Gq): 0.78±0.20 (n=5), p=0.34; **Fig. 11C, right**) and did not cause any significant change in the phosphorylation of pDARPP-32^S130^ (hM3D(Gq): 1.04±0.32 (n=6), p=0.89; **Fig. 11D, right**). These results showed that activation of Gq signaling cascades in D2-MSNS does not induce any of the changes in D1R expression and pDARPP-32phosphorylation of EAAC1^−/−^ mice.

If the cell-specific activation of Gq signaling cascades in D1-MSNs recapitulates the molecular phenotype of EAAC1^−/−^ mice, does it also reproduce the behavioral phenotype of EAAC1^−/−^ mice? To address this question, we repeated our grooming analysis in DREADD injected D1^Cre/+^ and A2A^Cre/+^ mice (**Fig. 11E,F**). As a control, to rule out potential off-target effects of CNO or its metabolites on endogenous receptors (Gomez et al., 2017), we performed stereotaxic injections of the DREADD construct or of saline solution and I/P CNO injections on age-matched WT mice (here referred to as Sham; **Fig. 11E,F**). Gq signaling activation caused an increase in the grooming frequency in D1^Cre/+^ mice versus Sham mice (Sham: 5.3±0.4e-3 Hz (n=25); D1^Cre/+^: 8.0±0.8e-3 Hz (n=39); D1^Cre/+^ vs Sham: **p=6.0e-3; **Fig. 11E,F**) without changing the duration of grooming episodes (Sham: 32.2±4.2 s (n=34); D1^Cre/+^: 40.9±3.5 s (n=44), p=0.12; **Fig. 11E,F**). Gq signaling activation in the DLS of A2A^Cre/+^ mice did not alter the grooming frequency and duration (6.0±0.4e-3 Hz (n=49), p=0.28; 42.3±3.9 s (n=58), p=0.08; **Fig. 11E,F**). The grooming behavior of D1^Cre/+^ mice that received DREADD activation was reminiscent of that of EAAC1^−/−^ mice, whereas the behavior of A2A^Cre/+^ mice was similar to that of WT mice in **Fig. 3A-B.** Therefore, cell-specific activation of Gq signaling cascades in D1-MSNs recapitulates the molecular and behavioral phenotype of EAAC1^−/−^ mice. These findings identify EAAC1 as a key regulator of glutamatergic and dopaminergic transmission and of repeated motor behaviors.

## DISCUSSION

Glutamate transporters play a fundamental role in shaping excitatory synaptic transmission in the brain. This knowledge largely relies on numerous and detailed studies of glial transporters, abundantly expressed in the neuropil (Danbolt, 2001). The neuronal glutamate transporter EAAC1 accounts for a small fraction of all glutamate transporters in the brain (Holmseth et al., 2012). As a consequence, its functional role has long remained enigmatic (Scimemi et al., 2009; Holmseth et al., 2012), despite evidence of genetic association between polymorphisms in the gene encoding EAAC1 and OCD (Dickel et al., 2006; Porton et al., 2013; Zike et al., 2017). In the hippocampus, EAAC1 limits extra-synaptic GluN activation by rapidly binding synaptically-released glutamate (Scimemi et al., 2009). The striatum is one of the brain regions where EAAC1 is most abundantly expressed (Danbolt, 2001; Kanai and Hediger, 2004), and is part of a poly-synaptic circuit that shows patterns of hyperactivity in patients with OCD (Ting and Feng, 2011; Ahmari et al., 2013). In the striatum, excitatory synaptic transmission is largely mediated by GluA receptors, with very little involvement of GluN receptors (Sung et al., 2001b). mGluRI glutamate receptors are abundantly expressed in the striatum (Albin et al., 1992; Shigemoto et al., 1992; Shigemoto et al., 1993; Testa et al., 1998; Ferraguti and Shigemoto, 2006) and their activation is crucial for the induction of long-term synaptic plasticity (Calabresi et al., 1992b; Anwyl, 1999; Sung et al., 2001a, b). A better understanding of EAAC1 in the context of OCD is important because small changes in the excitatory drive to striatal neurons can powerfully control the activity of neuronal circuits implicated with OCD (Radulescu et al., 2017).

Here we show that EAAC1 limits mGluRI activation in the striatum. By limiting mGluRI activation, EAAC1 promotes D1R expression and long-term plasticity. An interesting implication of our work is that intracellular Gq signaling cascades coupled to mGluRI activation are able to regulate both D1R expression and the phosphorylation state of DARPP-32 rapidly, within a 2-hour time window (**Fig. 11**). These findings are consistent with evidence that D1R internalization from the cell membrane can be modified within minutes using different pharmacological manipulations (Dumartin et al., 1998; Rassu et al., 2017).

An important insight of our work is that the functional implications of neuronal glutamate transporters differ across brain regions and should not rely exclusively on previous work in the hippocampus (Scimemi et al., 2009). Not only are there regional differences in the density of expression of EAAC1, but there are also regional differences in the distribution of other molecules susceptible to the activity of EAAC1 (e.g. GluN and mGluRI). The distinctive neuronal localization of EAAC1 puts it at an important positional advantage with respect to glial glutamate transporters, allowing it to be the first transporter to bind glutamate as it diffuses out of the synaptic cleft. It has been previously hypothesized that loss of EAAC1 may lead to increased glutamate concentration in the cerebrospinal fluid (Chakrabarty et al., 2005). Our experimental and modeling data suggest that this is an unlikely scenario, due to the low level of expression of EAAC1 (Holmseth et al., 2012). Instead, in the striatum and hippocampus, EAAC1 is implicated in the regulation of phasic synaptic transmission and long-term plasticity.

The findings that EAAC1^−/−^ mice show reduced D1R expression is consistent with recent radiolabeling assays, showing decreased D1R density in the DLS of mice with constitutively reduced EAAC1 expression via prolonged exposure to amphetamine (Zike et al., 2017). The DARPP-32 phosphoprotein is abundantly but not exclusively expressed in D1-MSNs (Ouimet et al., 1984; Anderson and Reiner, 1991) and is known to integrate glutamatergic and dopaminergic transmission in the striatum (Svenningsson et al., 2004). Our findings do not allow us to distinguish whether the effect of EAAC1 on D1R expression is directly mediated by DARPP-32 or by other molecules that are part of the Gq signaling cascades activated by mGluRI. However, they raise the possibility that these intracellular mechanisms may have different functional implications for integrating glutamatergic and dopaminergic signals in D1- and D2-MSNs.

Although many of the effects described in this work may appear subtle, they are noteworthy not only because they are statistically robust but because they yield novel insights into the physiological and pathological implications of EAAC1 for neuropsychiatric disorders that still lack effective pharmacological treatments. Loss of EAAC1 is associated with a distinctive behavioral phenotype which includes increased anxiety-like behaviors and disrupted grooming. Notably, the altered motor behaviors of EAAC1^−/−^ mice can be recapitulated by cell-specific activation of mGluRI-activated signaling cascades in D1-MSNs. The finding that increased mGluRI activation contributes to increased anxiety-like behaviors and repeated movement execution in EAAC1^−/−^ mice is consistent with a key role for these neuronal glutamate transporters in the modulation of motor function and perseverative behaviors. Our data are consistent with previous work in *Sapap3* knock-out mice, where increased mGluRI activation drives OCD-like behaviors via disruption of mGluR5-Homer interaction and constitutive mGluR5 signaling (Wan et al., 2011; Ade et al., 2016). Interestingly, PET studies in humans show that the mGluRI distribution volume ratio in brain regions that show functional abnormalities in OCD (e.g. cortico-striatal-thalamo-cortical pathway) is positively correlated with the patients' Y-BOCS obsession scores (Akkus et al., 2014). Increased expression and activation of mGluRI has also been observed in patients affected by other types of autism spectrum disorders (e.g. Fragile X syndrome, tuberous sclerosis complex) and intellectual disability (Boer et al., 2008; Maccarrone et al., 2010; Lohith et al., 2013), suggesting a possible common etiology for a broad range of neuropsychiatric diseases (D’Antoni et al., 2014). Many patients diagnosed with OCD are currently treated with cognitive behavioral therapy and/or serotonin reuptake inhibitors, but these approaches do not improve the symptoms of the disease for about 50% of the patients (Dougherty et al., 2004; Koran et al., 2007). One important implication of our work is that the treatment of neuropsychiatric disorders like OCD may be particularly challenging because it may require either cell-specific targeting of mGluRI or pharmacological manipulations affecting both glutamatergic and dopaminergic transmission in specific regions of the brain.

## ACKNOWLEDGEMENTS

This work was supported by SUNY Albany, SUNY Albany Research Foundation and the National Science Foundation (IOS 1655365). Thanks to Dimitri M. Kullmann and Marco Beato for comments on the manuscript.

## SUPPLEMENTARY INFORMATION

**Supplementary Figure 1-1. A battery of SHIRPA tests reveals subtle abnormalities in male and female EAAC1^−/−^ mice. (A)** Parameter scores for a battery of SHIRPA primary screen test in female WT (white bars) and EAAC1^−/−^ mice (red bars) aged P14-35. **(B)** As in (A), for male mice.

**Supplementary Figure 2-1. Female EAAC1^−/−^ mice show increased anxiety-like behaviors. (A)** Summary of spontaneous locomotor activity in a free-spinning flying saucer (left), in WT (n=60) and EAAC1^−/−^ female mice (n=64) aged P14-35. No significant difference was detected in the mean distance (left) and running time (right) between WT and EAAC1^−/−^ female mice (p=0.43 and p=0.26, respectively). (B) In the open field test, WT (n=66) and EAAC1^−/−^ female mice (n=59) travelled the same distance (p=0.14; left) but EAAC1^−/−^ female mice showed a trend to a decrease in the proportion of immobile time that did not reach statistical significance (p=0.078; right). **(C)** In the elevated plus maze (left), **EAAC1^−/−^** female mice (n=60) displayed a larger proportion of entries in closed arms in comparison to WT female mice (n=46; **p=0.001). The thick white lines represent the mean proportion of entries in each arm. **(D)** The proportion of time spent in the closed arms was also larger in EAAC1^−/−^ WT female mice (**p=0.002; right). **(E)** In the marble burial test (left), EAAC1^−/−^ mice (n=10; right) dug a significantly larger proportion of marbles than WT female mice (n=14, ***p=1.1e-7; left). **(F)** The color-coding represents the proportion of female mice digging a given proportion of marbles (y-axis). The white line describes the behavior of 50% of the mouse population. There is an average increase in the proportion of EAAC1^−/−^ female mice digging a larger proportion of marbles (*p=0.021).

**Supplementary Figure 2-2. Male EAAC1^−/−^ mice show increased anxiety-like behaviors. (A)** Summary of spontaneous locomotor activity in a free-spinning flying saucer (left), in WT *(n=77)* and EAAC1^−/−^ male mice (n=86) aged P14-35. No significant difference was detected in the mean distance (left) and running time (right) between WT and EAAC1^−/−^ male mice (p=0.56 and p=0.29, respectively). **(B)** In the open field test, WT (n=87) and EAAC1^−/−^ male mice (n=90) travelled the same distance (p=0.90; left) but EAAC1^−/−^ male mice showed a decreased the proportion of immobile time (*p=0.033; right). **(C)** In the elevated plus maze, EAAC1^−/−^ male mice (n=30) displayed an increased proportion of entries in the closed arms in comparison to WT male mice (n=72, *p=0.046). The thick white lines represent the mean proportion of entries in each arm. **(D)** The proportion of time spent in the closed arms was also larger in EAAC1^−/−^ WT male mice (*p=0.036; right). **(E)** In the marble burial test (left), EAAC1^−/−^ mice (n=5; right) dug a significantly larger proportion of marbles than WT male mice (n=22, **p=2.5e-9; left). **(F)** The color-coding represents the proportion of male mice digging a given proportion of marbles (y-axis). The white line describes the behavior of 50% of the mouse population. There is an average increase in the proportion of EAAC1^−/−^ male mice digging a larger proportion of marbles (*p=0.049).

**Supplementary Figure 7-1. Validation of D1R and D2R antibody specificity. (A)** Left: Confocal images of striatal sections from the DLS of D1^Cre/+^:Ai9^Tg/0^ (top panel) and A2A^Cre/+^:Ai9^Tg/0^ mice (bottom panel), immuno-labelled for D1R (green). D1- and D2-MSNs express the red fluorescent protein mCherry in D1^Cre/+^:Ai9^Tg/0^ and A2A^Cre/+^:Ai9^Tg/0^ mice, respectively. Right: quantification of co-labelled red (R), green (G) and yellow cells (Y). The Y/R ratio in D1^Cre/+^:Ai9^Tg/0^ (green bar) and A2A^Cre/+^:Ai9^Tg/0^ mice (red bar) represents the proportion of mCherry-expressing cells recognized by the anti-D1R antibody. The Y/R ratio is large in D1^Cre/+^:Ai9^Tg/0^ mice (n=4) and small in A2A^Cre/+^:Ai9^Tg/0^ mice (n=3). The Y/G ratio in D1^Cre/+^:Ai9^Tg/0^ (green bar) and A2A^Cre/+^:Ai9^Tg/0^ mice (red bar) represents the proportion of all immuno-labelled cells recognized by the anti-D1R antibody that also express mCherry. The Y/G ratio is large in D1^Cre/+^:Ai9^Tg/0^ mice (n=4) and small in A2A^Cre/+^:Ai9^Tg/0^ mice (n=3). **(B)** As in (A), for the anti-D2R antibody. The Y/R ratio in D1^Cre/+^:Ai9^Tg/0^ (green bar) and A2A^Cre/+^:Ai9^Tg/0^ mice (red bar) represents the proportion of mCherry-expressing cells recognized by the anti-D2R antibody. The Y/R ratio is small in D1^Cre/+^:Ai9^Tg/0^ mice (n=3) and large in A2A^Cre/+^:Ai9^Tg/0^ mice (n=4). The Y/G ratio is small in D1^Cre/+^:Ai9^Tg/0^ mice (n=3) and large in A2A^Cre/+^:Ai9^Tg/0^ mice (n=4). **(C)** Western blot analysis of anti-D1R antibody specificity. D1R expression was significantly lower in the cortex compared to the striatum (Norm D1R striatum (n=8), Norm D1R cortex (n=8), *p=0.037). No band was detected when the antibody was pre-adsorbed with the D1R peptide control (Norm D1R peptide (n=7), ***p=3.7e-4). **(D)** As in (C), for anti-D2R antibody. D2R expression in the cortex was significantly lower than in the striatum (Norm D2R striatum (n=14), Norm D2R cortex (n=16), *p=0.022). No band was detected by the anti-D2R antibody in pre-adsorption experiments (Norm D2R peptide (n=7), ***p=1.1e-6).

**Supplementary Figure 7-2. Reduced D1R expression in EAAC1^−/−^ mice is not due to fewer D1-MSNs. (A)** Confocal images of sections from D1^Cre/+^: Ai9^Tg/0^ and D1^Cre/+^: Ai9^Tg/0^: EAAC1^−/−^ mice (red (R): mCherry; blue (B): DAPI). **(B)** Top: summary graph showing similar density of D1-MSNsin the DLS (WT (n=ll), EAAC1^−/−^ (n=6), p=0.90) and VMS of WT and EAAC1^−/−^ mice (WT (n=6), EAAC1^−/−^ (n=6), p=0.073). Bottom: there is a similar proportion of D1-MSNs in the DLS (WT (n=11), EAAC1^−/−^ (n=6), p=0.64) and VMS of WT and EAAC1^−/−^ mice (WT (n=7), EAAC1^−/−^ (n=6), p=0.055). **(C)** Confocal images of sections from A2A^Cre/+^: Ai9^Tg/0^ and A2A^Cre/+^: Ai9^Tg/0^: EAAC1^−/−^ mice. Top: summary graph showing similar density of D2-MSNsin the DLS (WT (n=6), EAAC1^−/−^ (n=5), p=0.14) and VMS of WT and EAAC1^−/−^ mice (WT (n=6), EAAC1^−/−^ (n=5), p=0.58). The density of D2-MSNs is larger in the DLS than in the VMS for both WT (**p=1.7e-4) and EAAC1^−/−^ mice (*p=0.022). Bottom: there is a similar proportion of D2-MSNs in the DLS (WT (n=6), EAAC1^−/−^ (n=5), p=0.93) and VMS of WT and EAAC1^−/−^ mice (WT (n=6), EAAC1^−/−^ (n=5), p=0.31).

